# A complex role of Arabidopsis CDKD;3 in meiotic progression and cytokinesis

**DOI:** 10.1101/2022.08.08.503215

**Authors:** Sorin Tanasa, Neha Shukla, Albert Cairo, Ranjani S. Ganji, Pavlina Mikulková, Sona Valuchova, Vivek K. Raxwal, Claudio Capitao, Arp Schnittger, Zbyněk Zdráhal, Karel Riha

## Abstract

Meiosis is a specialized cell division that halves the number of chromosomes in two consecutive rounds of chromosome segregation. In angiosperm plants is meiosis followed by mitotic divisions to form rudimentary haploid gametophytes. In Arabidopsis, termination of meiosis and transition to gametophytic development is governed by TDM1 and SMG7 that mediate inhibition of translation. Mutants deficient in this mechanism do not form tetrads, but instead undergo multiple cycles of aberrant nuclear divisions, that are likely caused by the failure to downregulate cyclin dependent kinases during meiotic exit. A suppressor screen to identify genes that contribute to meiotic exit uncovered a mutation in CDKD;3 that alleviates meiotic defects in *smg7* deficient plants. The CDKD;3 deficiency prevents aberrant meiotic divisions observed in *smg7* mutants, or delays their onset after initiation of cytokinesis, which permits formation of functional microspores. Although CDKD;3 acts as an activator of CDKA;1, the main cyclin dependent kinase that regulates meiosis, *cdkd;3* mutation appears to promote meiotic exit independently of CDKA;1. Furthermore, analysis of CDKD;3 interactome revealed enrichment for proteins implicated in cytokinesis suggesting a more complex function of CDKD;3 in cell cycle regulation.

## Introduction

The eukaryotic cell cycle consists of two major cellular events: DNA replication in S-phase and chromosome segregation followed by cell division in M-phase. Alteration of these events is accomplished by executing complex molecular processes that take place in a well-defined order to ensure faultless passage of genetic information from one cell generation to another. Cyclin-dependent kinases (CDKs) are the master regulators of the eukaryotic cell cycle. CDKs push cells into S-phase through activation of S-phase specific genes and initiation of DNA replication. Further buildup of CDK activity at G2/M drives chromosome condensation, dissolution of the nuclear membrane, formation of the mitotic spindle and other processes required for chromosome segregation and cell division. CDK activity peaks at metaphase when all chromosomes properly attach to the spindle microtubules. This triggers a rapid decline of CDK activity and the dephosphorylation of its targets, which allows for chromosome decondensation, spindle disassembly and cytokinesis. Low CDK activity in G1 is crucial for DNA replication licensing, which assures that chromosomes are duplicated only once during the cell cycle.

The plant cell cycle is driven by two types of CDKs: the canonical CDKA corresponding to the mammalian CDK1 is expressed throughout the cell cycle, and the plant-specific CDKBs whose expression is cell-cycle dependent (Shimotohno *et al*., 2021). CDK activity and specificity are controlled by cyclins, which in *Arabidopsis thaliana* form a large gene family with at least 50 members (Vandepoele *et al*., 2002, Wang *et al*., 2004a). Type A, B and D cyclins are directly implicated in cell-cycle and their consecutive activities drive progression through S-, G2- and M-phases. Cyclin levels oscillate due to ubiquitin-mediated degradation resulting in waves of CDK activities as the cell cycle progresses.

CDKA and CDKBs are further regulated by phosphorylation in their T-loop by CDK-activating kinases that consist of CDKDs and cyclin H (Yamaguchi *et al*., 1998, Umeda *et al*., 2005, Dissmeyer *et al*., 2007, Harashima *et al*., 2007). Plant CDKDs correspond to mammalian CDK7; like CDKD7, they are capable of not only activating other CDKS, but also phosphorylating the C-terminal domain of the largest subunit of RNA polymerase II, thereby affecting transcription and co-transcriptional RNA processing (Yamaguchi *et al*., 1998, Shimotohno *et al*., 2003, Hajheidari *et al*., 2012). Arabidopsis encodes three CDKD paralogues, CDKD;1, CDKD;2 and CDKD;3 that are partially redundant and their concomitant inactivation is lethal (Hajheidari *et al*., 2012, Takatsuka *et al*., 2015). Further functional studies showed that CDKDs regulate microtubule dynamic and meiotic cytokinesis through activation of CDKA;1, and small RNA biogenesis through phosphorylation of the CTD domain of RNA polymerase II (Hajheidari *et al*., 2012, Sofroni *et al*., 2020).

Meiosis is a specialized form of cell division that produces haploid cells from a diploid progenitor in two successive rounds of chromosome segregation without intervening S-phase. The meiotic program substantially deviates from regular mitotic division and its implementation requires extensive modification of chromosome segregation and cell-cycle machinery. These include physical coupling of homologous chromosomes via recombination and synapsis in prolonged prophase I, mono-orientation of sister kinetochores in metaphase I, protection of centromeric cohesion during anaphase I, and inhibition of DNA replication in interkinesis. These processes are brought about by specific sets of effector proteins and cell-cycle regulators, many of which are uniquely expressed in meiosis. Despite the differences between mitosis and meiosis, the core molecular machinery driving their progression seems to be identical. For example, the key Arabidopsis mitotic kinase, CDKA;1, appears to be central also for meiosis where it orchestrates chromosome recombination and pairing, spindle organization and cytokinesis (Wijnker *et al*., 2019, Yang *et al*., 2020, Sofroni *et al*., 2020). This is done in concert with several cyclins, of which some are meiosis specific, while others are present also in mitosis (Azumi *et al*., 2002, Wang *et al*., 2004b, Bulankova *et al*., 2013).

In angiosperm plants, haploid products of meiosis undergo additional mitoses to generate rudimentary haploid gametophytes that carry germ cells. Therefore, upon completion of meiosis, the cell division machinery is reprogrammed back to mitosis. Male germ cells differentiate in pollen, a tricellular structure produced by two consecutive post-meiotic divisions of a haploid microspore. Studies in Arabidopsis revealed a dedicated molecular mechanism that governs the termination of male meiosis and transition to post-meiotic development. This mechanism relies on the evolutionarily conserved nonsense-mediated RNA decay (NMD) factor SMG7 and plant specific protein TDM1, and facilitates meiotic exit through repressing translation (Cairo *et al*., 2022). Mutants deficient in these proteins fail to terminate meiosis and to produce pollen. In *tdm1* mutants, meiosis progresses normally until telophase II, but instead of undergoing cytokinesis, the haploid nuclei attempt to further divide without replicating their DNA. Such pollen mother cells (PMCs) enter multiple cycles of chromatin condensation and spindle re-assembly, which leads to unequal distribution of chromatin and formation of polyads (Bulankova *et al*., 2010, Cairo *et al*., 2022). A similar phenotype was caused by partially functional allele *smg7-6*, whereas the full SMG7 inactivation halts meiotic progression in anaphase II (Riehs *et al*., 2008, Capitao *et al*., 2021). Such phenotypes are reminiscent of situations when cells fail to downregulate CDK activity at the end of M-phase (Parry and O’Farrell, 2001, Potapova *et al*., 2006). Indeed, *smg7* mutants arrested at anaphase II exhibit high level of T-loop phosphorylated CDKA;1 indicating that high CDK activity persists beyond metaphase II (Bulankova *et al*., 2010).

To decipher molecular processes that govern meiotic exit, we performed a forward genetic screen to identify mutations that suppress phenotype of *smg7-6* plants and increase their fertility. We anticipated to recover mutations either directly affecting the TDM1/SMG7-mediated mechanism that regulates meiotic exit, or rescuing the fertility indirectly through modulation of chromosome segregation or cell cycle progression. In this study we report on characterizing suppressor mutation in CDKD;3 and its effect on meiotic progression and exit in *smg7-6* plants.

## Results

### Characterization of EMS58-1 suppressor line exhibiting recovered fertility

Arabidopsis mutants homozygous for the hypomorphic *smg7-6* allele have reduced fertility due to 10-fold decrease of viable pollen compared to wild type (Figure 1) (Riehs-Kearnan *et al*., 2012, Capitao *et al*., 2021). A characteristic feature of *smg7-6* plants is that the first 15-20 flowers on the main inflorescence bolt are sterile, but later flowers regain fertility. To identify *smg7-6* suppressors, we mutagenized the seeds homozygous for *smg7-6* with ethyl-methanesulfonate (EMS) and screened for plants with increased length of the first 20 siliques in the M2 generation, which was indicative of restored seed production (Capitao *et al*., 2021). One of the suppressor lines named *EMS58-1* exhibited increased fertility and about 4-fold increase in viable pollen compared to *smg7-6* parents. However, the pollen count was still lower than in wild type (Figure 1). The *EMS58-1* line did not show any obvious growth abnormalities and, with the exception of increased fertility, it was indistinguishable from *smg7-6* parents (Figure S1).

**Figure 1.**
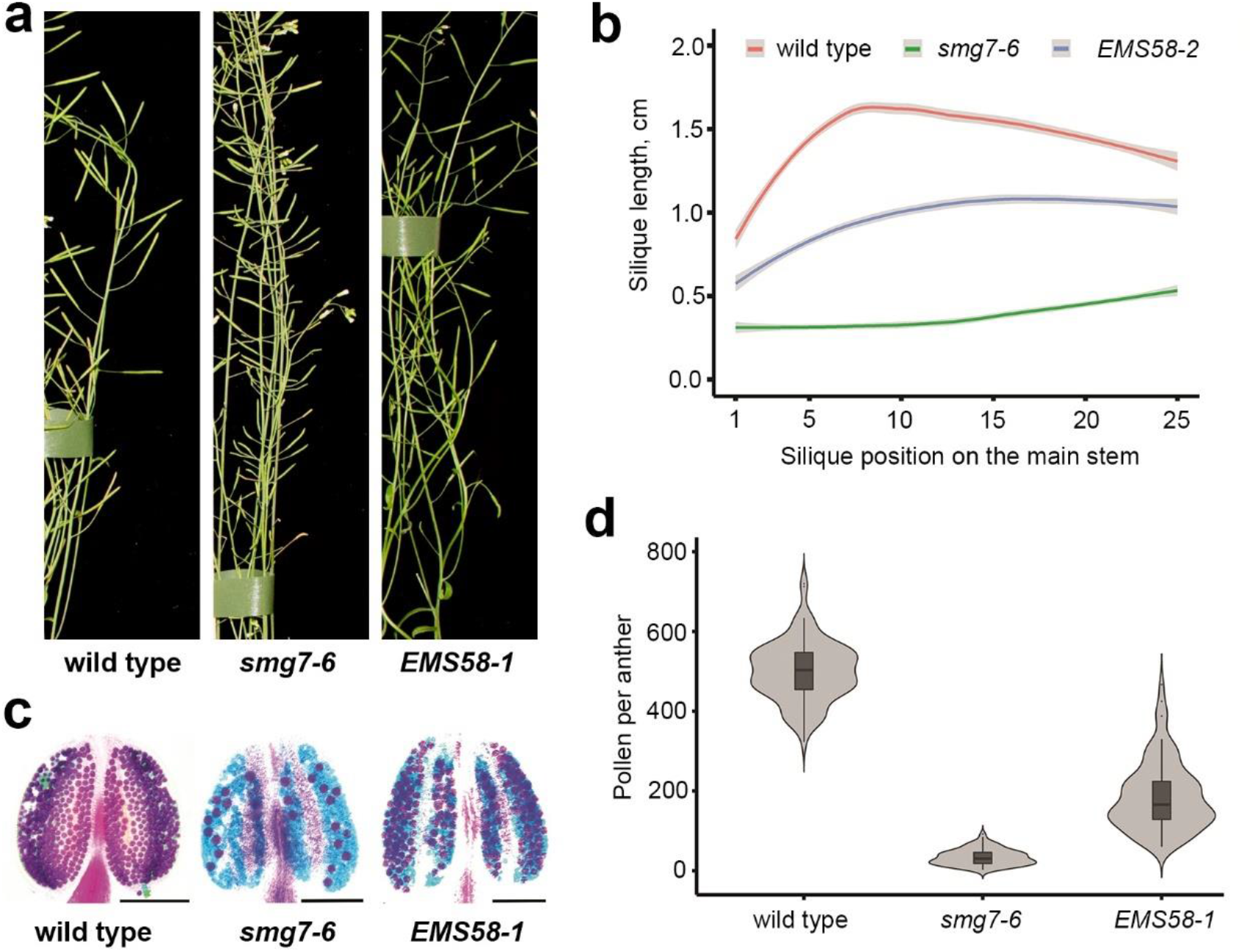
Fertility of the EMS58-1 suppressor line. (a) Details of inflorescence bolt with siliques from 6 weeks old plants depicted in Figure S1. (b) Quantification of silique length along the main inflorescence bolt in wild type (n=25), *smg7-6* (n=37) and *EMS58-2* (n=48). The trend lines of the data were plotted by LOESS smooth function (colored lines; shaded area represents 95% confidence intervals) (c) Anthers of indicated lines after Alexander staining. Scale bar = 10 µm. (d) Violin plots showing viable pollen per anther (wild type n= 96; *smg7-6* n=97; *EMS58-1* n=97).

PMCs in *smg7-6* mutants do not conclude meiosis by cytokinesis. Instead, haploid nuclei undergo multiple rounds of chromatin condensation/decondensation and spindle assembly/reassembly (referred here to as meiosis III, meiosis IV etc.; (Capitao *et al*., 2021). This results in random distribution of chromatids and formation of polyads in more than 95% of PMCs (Figure 2a,b). In contrast, we observed that 20 - 80% of PMCs in individual anther lobes in EMS58-1 plants formed tetrads. This result was further reaffirmed by crossing *EMS58-1* with *qrt1* plants that are deficient in pectin methylesterase and do not separate pollen derived from the same PMC (Francis *et al*., 2006). The fraction of tetrads containing four viable pollen rose from 21% in *smg7-6 qrt1* to 52% in *EMS58-1 qrt1* pants (Figure 2c). These data suggest that a substantial portion of PMCs in *EMS58-1* properly terminate meiosis and do not enter aberrant postmeiotic chromosome segregation.

**Figure 2.**
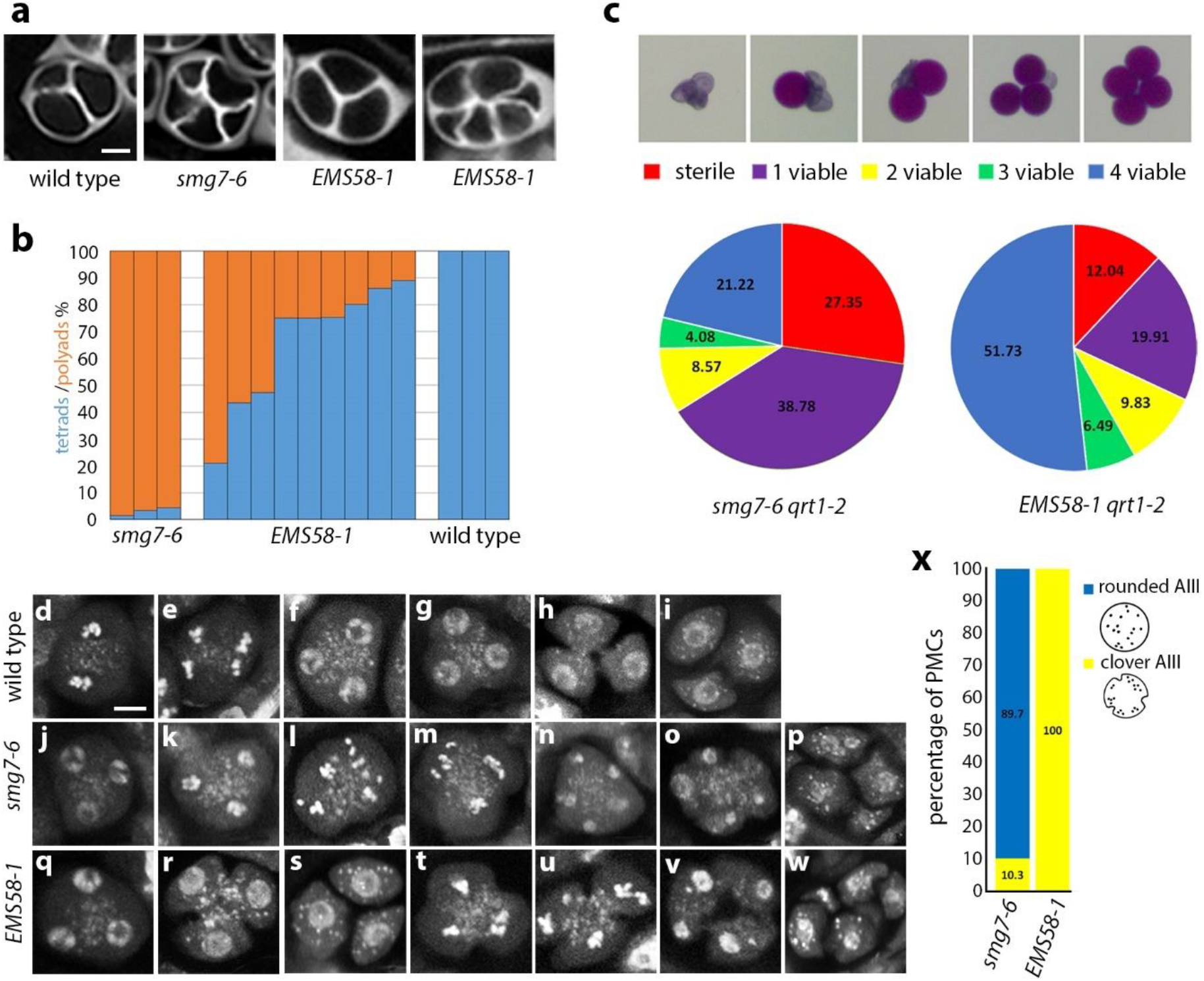
Male meiosis in the EMS58-1 suppressor line. (a) Tetrads and polyads in indicated mutants visualized by callose staining with SCRI Renaissance 2200 (SR2200). Scale bar = 5 µm. (b) Frequency of tetrads and polyads in individual anthers. (c) Viability of pollen derived from the same tetrad assessed by Alexander staining. Pie charts show proportion of tetrads with indicated number of viable pollens in *smg7-6 qrt1-2* (n = 245 tetrads) and *EMS58-1 qrt1-2* (n = 1219). (d-w) PMCs and tetrads stained by DAPI. Figures (k-p) and (t-w) show aberrant post-meiotic divisions resulting in polyads, whereas (g-h) and (q-s) represent regular cytokinesis and tetrad formation Scale bar = 5 µm; (d) metaphase II, (e) anaphase II, (m,t,u) clover anaphase III, (l) rounded anaphase III. (x) Frequency of rounded and clover anaphases III in smg7-6 (n = 183) and *EMS58-1* mutants (n = 74).

Interestingly, we also noticed that a portion of tetrads harboring two or three viable pollen is higher relative to tetrads with no or a single pollen in *EMS58-1 qrt1* compared to *smg7-6 qrt1* (Figure 2c). We assume that PMCs producing 1 to 3 viable pollen represent situations where cells entered aberrant postmeiotic segregation, but the original set(s) of chromosomes did not fully separate and remained clustered forming functional haploid nuclei (Capitao *et al*., 2021). A higher frequency of tetrads with 2 to 3 viable pollen in line *EMS58-1* indicated increase propensity of chromosomes to stay together during aberrant divisions.

Cytogenetic analysis showed two populations of PMCs in *EMS58-1* plants: ones that terminated meiosis normally and formed tetrads (Figures 2d-i and 2q-s), and the other ones that entered aberrant chromosome segregation (Figures 2t-w) typical for *smg7-6* mutants (Figures 2j-p). However, there was a difference between *smg7-6* and *EMS58-1* PMCs undergoing aberrant segregations. Whereas *smg7-6* PMCs progressing through meiosis III were mostly rounded (Figure 2l,x), the majority of *EMS58-1* meiocytes in anaphase III were pinched inwards as if undergoing cytokinesis (referred to as “clover” PMCs; Figures 2t,u and x). This observation indicated that the onset of cytokinesis precedes the completion of aberrant chromosome segregation in *EMS58-1* plants.

We next performed live cell imaging of PMCs to assess meiotic progression in *EMS58-1*. We used two experimental setups: in the first one, we measured the duration of stages when chromatin is condensed (meiosis I, meiosis II, meiosis III) and decondensed (interkinesis I and interkinesis II) using HTA10:TagRFP chromatin marker (Figure 3a, Movies 1-3). In the second setup, we measured the time interval between spindle disassembly either in meiosis II or meiosis III and cytokinesis using the TagRFP:TUB4 reporter for microtubules (Figure 3b, Movies 4-6). Together, these markers allowed us to assess the duration of meiosis from diakinesis/metaphase I to cytokinesis. *EMS58-1* meiocytes that formed regular tetrads were compared to wild type, whereas *EMS58-1* meiocytes undergoing meiosis III were compared to *smg7-6*. This analysis revealed that all stages of meiosis were delayed in *EMS58-1* plants compared to the respective controls (Figure 3c-e). Nevertheless, the most pronounced difference was observed in interkinesis II and cytokinesis in PMCs undergoing meiosis III, both of which lasted ∼ 1 h longer in *EMS58-1* than in *smg7-6* (Figure 3e). We further noticed that chromatids in *EMS58-1* PMCs undergoing anaphase III tend to spread less than chromatids in *smg7-6* (Figure S2).

**Figure 3.**
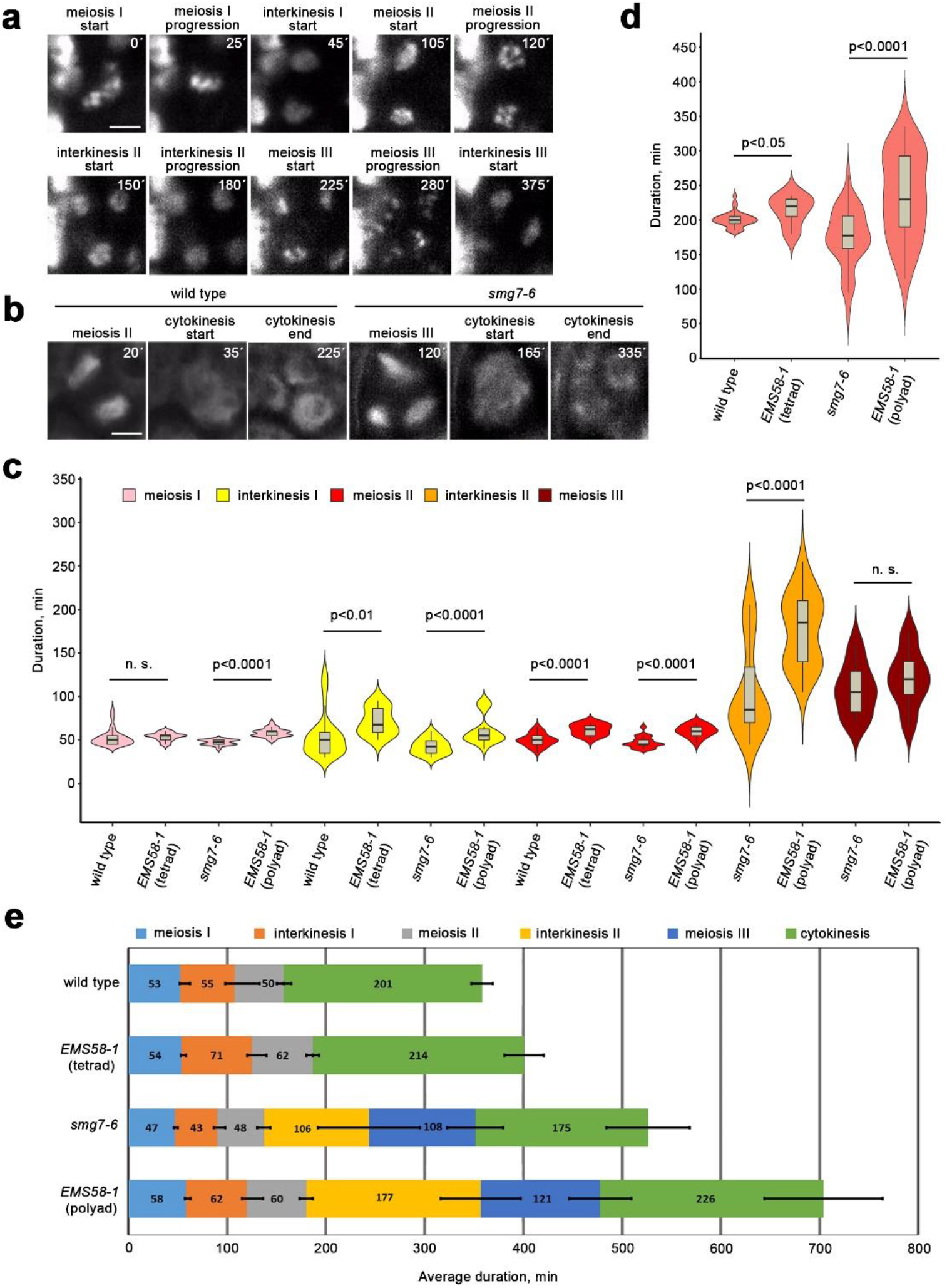
Live imaging of meiotic progression in *EMS58-1*. (a) Time-lapse series of *smg7-6* meiocyte with HTA10:TagRFP labeled chromatin. Time points relate to metaphase I. Scale bar = 5 µm. (b) Time-lapse series of wild type and *smg7-6* meiocytes containing microtubules labeled with TagRFP:TUB4 reporter. Time points relate to the formation of meiosis I spindle. Scale bar = 5 µm. (c) Violin plots showing duration of individual meiotic stages assessed from live cell imaging of PMCs with labeled chromatin. Significance of the difference is indicated (two tailed t-test; wild type n = 25, *EMS58-1* (tetrad) n = 16, *smg7-6* n = 18, *EMS58-1* (polyad), n = 23). (d) Violin plots showing duration of cytokinesis assessed from live cell imaging of PMCs with labeled microtubules. Significance of the difference is indicated (two tailed t-test; wild type n=35, *EMS58-1* (tetrad) n=5, *smg7-6* n = 40, *EMS58-1* (polyad) n = 35). (e) Graphical representation of meiotic duration based on data from (c) and (d). Error bars represent SDs for the duration of each meiotic stage.

Together, these data indicate two mechanisms that contribute to the increased fertility and pollen production in the *EMS58-1* line. First, a portion of meiocytes does not enter aberrant postmeiotic divisions and forms normal tetrads after meiosis II. In the remaining PMCs is meiosis III substantially delayed and coincides with the onset of cytokinesis, which restricts the movement of chromatids and increases the chance for assembly of haploid nuclei with the full set of chromosomes required for pollen formation.

### Restored fertility is caused by mutation in CDKD;3

We used association mapping to identify the mutation restoring fertility in the *EMS58-1* line. We backcrossed *EMS58-1* to the parental *smg7-6* line and established the B2 mapping population. The increased fertility phenotype segregated as a recessive trait. The associated *de novo* mutations were identified by comparative whole genome sequencing of fertile B2 plants with *smg7-6* parents. This analysis revealed *de novo* mutations that associated with fertility on chromosome 1 between 5.5 and 7 Mb (Figure S3, Table S1). PCR-based genotyping of several selected mutations in this region in 106 B2 plants showed the highest association between phenotype and mutation in the At1g18040 locus coding for CDKD;3. The mutation represents a C to T transition in the third exon of the gene and causes a proline to leucine substitution (P137L) at amino acid position 137 (Figure 4a). The P137L mutation is located in the evolutionary highly conserved kinase active site and likely impairs CDKD;3 activity (Figure 4b). We refer to this new allele as *cdkd;3-3*. Transformation of *EMS58-1* plants with a wild copy of the CDKD;3 gene reduced fertility in the majority of T1 transformants back to the level observed in *smg7-6*, confirming the causality of the mutation (Figure S4). When introgressed to wild type Col-0 background, *cdkd;3-3* mutants did not exhibit any obvious phenotype and were fully fertile (Figure S5).

**Figure 4.**
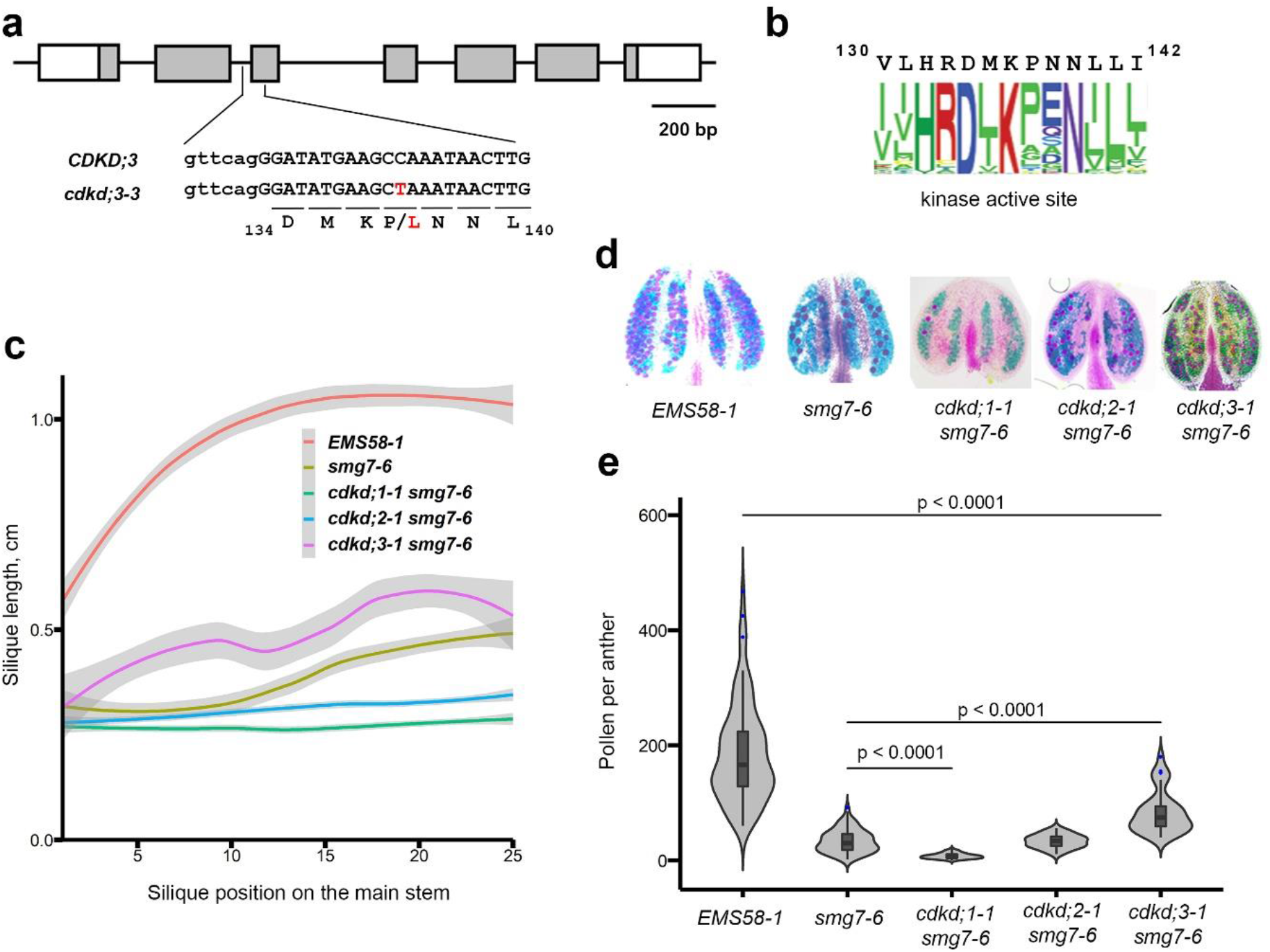
Effect of CDKDs on fertility of smg7-6 plants. (a) Diagram *of CDKD;3* gene with exons marked as boxes. DNA sequence surrounding the *de novo* mutation (depicted in red) in the *cdkd;3-3* allele is shown; capital letters denote exon sequence. (b) The conserved amino acid sequence present in the kinase active site of CDKD3 is represented as web logo. A total of 131,130 kinase sequences were aligned with CDKD3 active kinase site (amino acid residues 130 to 142). (c) Quantification of silique length along the main inflorescence bolt in wild type and indicated mutants (*smg7-6* n = 25, *cdkd;1-1 smg7-6* n = 20, *cdkd;2-1 smg7-6* n = 20, *cdkd;3-1 smg7-6* n = 11). The trend lines of the data were plotted by LOESS smooth function (colored lines; shaded area represents 95% confidence intervals). (d) Anthers of indicated mutants after Alexander staining. (d) Violin plots showing viable pollen per anther. Significance of the difference is indicated (two tailed t-test; *EMS58-1* - 97 anthers/10 plants, *smg7-6* - 94 anthers/10 plants, *cdkd;1-1 smg7-6* - 70 anthers/6 plants, *cdkd;2-1 smg7-6* - 35 anthers/3 plants, *cdkd;3-1 smg7-6* - 36 anthers in 4 plants).

Because Arabidopsis CDKD;3 paralogues exhibit functional redundancy (Hajheidari *et al*., 2012, Takatsuka *et al*., 2015), we next examined the impact of each paralogue on the fertility of *smg7-6* plants. We crossed T-DNA insertion alleles *cdkd;1-1, cdkd;2-1* and *cdkd;3-1* individually with *smg7-6* and examined fertility in the respective double mutants (Figure S6). Only *cdkd;3-1 smg7-6* plants showed partially improved fertility and pollen count (Figure 4c-e). Inactivation of the other two CDKDs further decreased fertility of *smg7-6*, with *cdkd;1-1 smg7-6* plants producing no pollen. This data suggests partial functional divergence of the Arabidopsis CDKD paralogues. Interestingly, the *cdkd;3-1* allele improved pollen yield to a much lesser extent than *cdkd;3-3* allele (Figure 4f). We suspect that the CDKD;3^P137L^ variant produced from *cdkd;3-3* sequesters cyclin H and other subunits in a non-productive complex, therefore exerting a bigger effect on CDKD activity than full disruption of the gene in *cdkd;3-1* plants.

### Downregulation of CDKA;1 does not restore meiotic exit in smg7-6 mutants

Arabidopsis CDKDs were described as the CDKA;1 activating kinases and CDKA;1 appears to be a key driver of meiotic progression. In Arabidopsis, cytokinesis occurs simultaneously after meiosis is completed (De Storme and Geelen, 2013). Inhibition of cytokinesis after meiosis I is mediated by CDKA;1 and combined reduction of CDKD; and CDKA;1 activities led to sequential cytokinesis or premature meiotic exit after meiosis I, depending on the extent of the reduction (Sofroni *et al*., 2020). Our previous work hinted that aberrant meiotic exit in SMG7 deficient plants may be caused by the failure to downregulate CDKA;1 (Bulankova *et al*., 2010). Therefore, identification of CDKD;3 in the suppressor screen is consistent with the scenario where CDKD insufficiency leads to a lower level of CDKA;1, which in turn promotes cytokinesis and meiotic exit in *smg7-6* plants.

To directly test this scenario, we combined *smg7-6* mutation with a line where endogenous CDKA;1 was substituted with its hypomorphic CDKA;1^T14V;Y15F^ variant (CDKA;1^VF^) that exhibits lower activity (Dissmeyer *et al*., 2009). The *CDKA;1*^*VF*^ plants have partially reduced fertility and pollen count (Figure 5a-c), which is consistent with the key role of CDKA;1 in meiotic progression. Cytogenetic analysis showed that many CDKA;1^VF^ PMCs initiate cell wall formation prior to the tetrad stage (Figure 5d-f). This leads to partial separation of interkinesis nuclei, which permits more flexibility in spindle orientation during meiosis II, resulting in tetrad configurations reflecting tetrahedral (regular), parallel or perpendicular divisions (Figure 5d-f). Surprisingly, CDKA;1^VF^ did not suppress the *smg7-6* phenotype, and meiocytes in *CDKA;1*^*VF*^ *smg7-6* plants progressed into aberrant post-meiotic divisions and formed polyads (Figure 5d-f). Furthermore, *smg7-6* prevented the partial cytokinesis observed in *CDKA;1*^*VF*^ plants (Figure 5e). These data indicate that suppression of aberrant meiosis and promotion of cytokinesis in EMS58-1 plants is mediated through a CDKA;1 independent mechanism.

**Figure 5.**
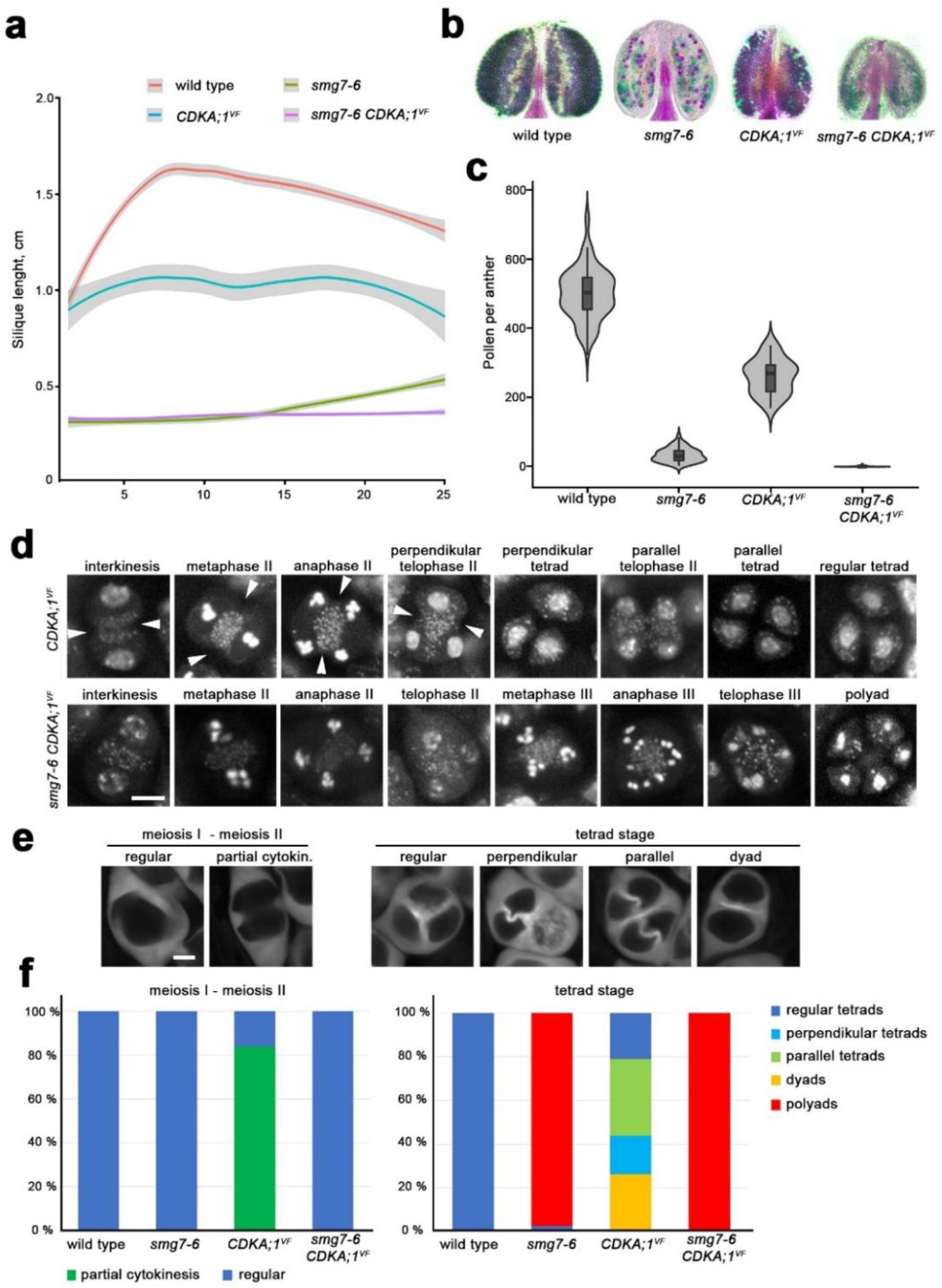
Genetic interaction between *CDKA;1*^*VF*^ and *smg7-6*. (a) Quantification of silique length along the main inflorescence bolt in wild type and indicated mutants (wild type n = 25, *smg7-6* n = 25, *CDKA;1*^*VF*^ n = 9 and *CDKA;1*^*VF*^ *smg7-6* n = 23 plants). The trend lines of the data were plotted by LOESS smooth function (colored lines; shaded area represents 95% confidence intervals). (b) Anthers of indicated mutants after Alexander staining. (c) Violin plots showing viable pollen per anther (wild type n =96, *smg7-6* n = 94, *CDKA;1*^*VF*^ n = 30 and *CDKA;1*^*VF*^ *smg7-6* n = 116). (d) Cytogenetic analysis of meiosis II and cytokinesis in DAPI stained PMCs. Arrowheads point to cell wall invaginations prior to completion of meiosis II. Scale bar = 5 µm. (e) Callose staining of *CDKA;1*^*VF*^ PMCs with SR2200 dye to visualize cell wall. Scale bar = 5 µm. (f) Quantification of PMCs undergoing partial cytokinesis prior to completion of chromosome segregation (left panel; wild type n = 86, *smg7-6* n = 166, *CDKD;1*^*VF*^ n = 155, *smg7-6 CDKD;1*^*VF*^ n = 95) and tetrad configurations after cytokinesis (right panel; wild type n = 200, *smg7-6* n = 524, *CDKD;1*^*VF*^ n = 256, *smg7-6 CDKD;1*^*VF*^ n = 246).

To further explore the role of CDKD;3 in meiotic cytokinesis, we analyzed its genetic interaction with *TAM* (Wang *et al*., 2004b). TAM is an A-type cyclin expressed during meiosis I whose inactivation results in meiotic exit after meiosis I and formation of diploid microspores. Due to the absence of meiosis II, the pollen count in *tam* mutants is reduced to half compared to wild type (Figure 6a,b). Live cell imaging revealed that *tam* mutants also form ectopic spindle-like and phragmoplast-like structures in the cytoplasm of prophase I PMCs (Prusicki *et al*., 2019). Consistent with these data, we observed that PMCs in TAM deficient plants are often pinched and form bud-like structures that indicate attempts of ectopic and premature cytokinesis (Figure 6c,d). The ectopic cytokinesis was substantially reduced in *cdkd;3-3 tam* double mutants (Figure 6e). Thus, the effect of *cdkd;3-3* mutation on cytokinesis is context-dependent: while it reduces the premature cytokinesis during prophase I in TAM-null plants, it promotes cytokinesis after meiosis II in *smg7-6* mutants.

**Figure 6.**
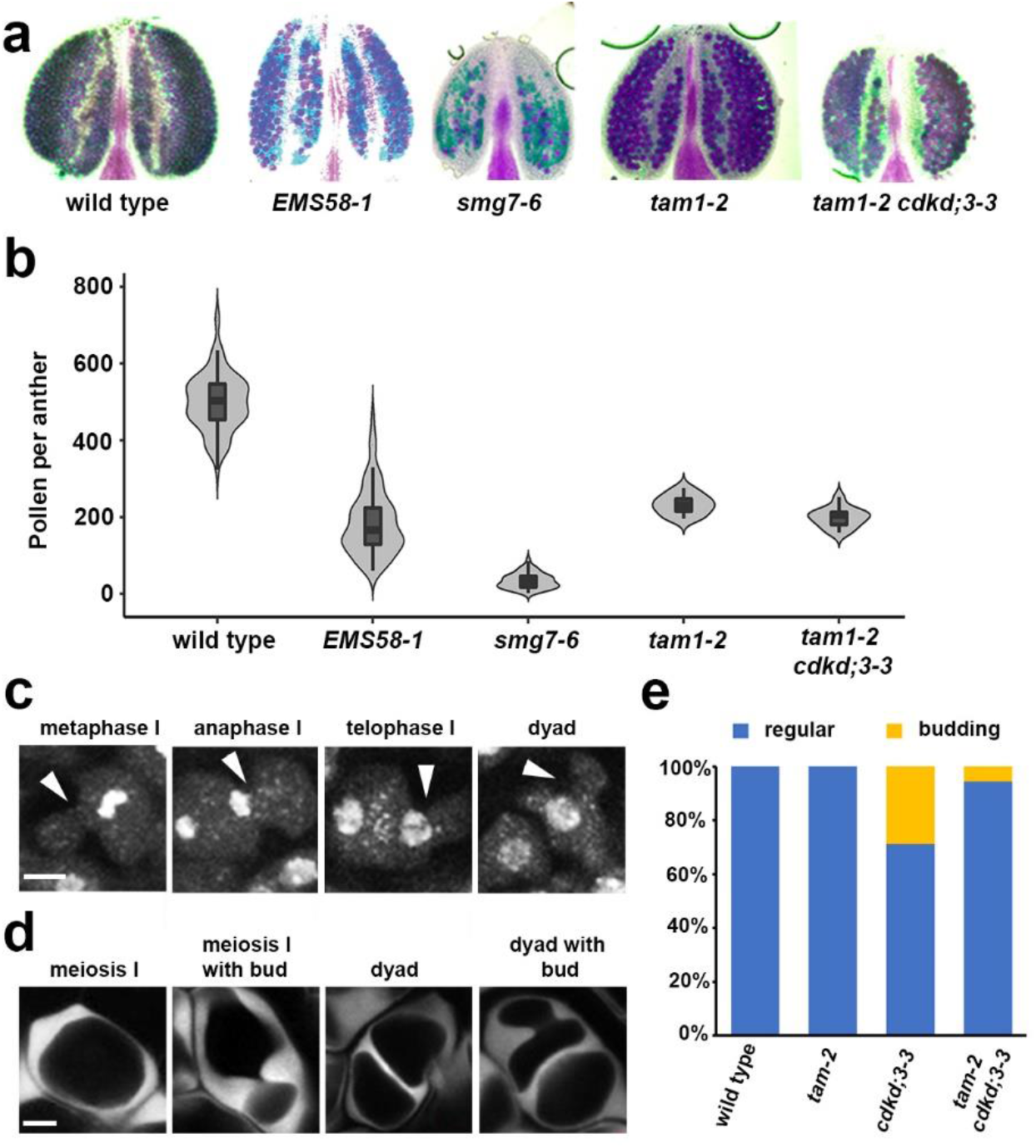
Genetic interaction between *tam* and *cdkd;3-3*. (a) Anthers of indicated mutants after Alexander staining. (c) Violin plots showing viable pollen per anther (wild type n = 96, *EMS58-1* n = 97, *smg7-6* n = 94, *tam-2* n = 10, *tam-2 cdkd;3-3* n = 13). (c) Cytogenetic analysis of meiosis I in DAPI stained PMCs of *tam*. Arrowheads point to ectopic cell wall invaginations that form buds during metaphase I. Scale bar = 5 µm. (e) Callose staining of *tam* PMCs with SR2200 dye to visualize cell wall. Scale bar = 5 µm. (f) Quantification of PMCs exhibiting ectopic cell wall formation and budding. (wild type n = 386, *cdkd;3-3* –n =455, *tam-2* n =187 and *tam-2 cdkd;3-3* n = 101).

### Exploration of alternative mechanisms of CDKD;3 action by transcriptomics and proteomics

Our data indicate that *cdkd;3-3* may restore the fertility of *smg7-6* plants independently of the CDKD function in activating CDKA;1. The CTD domain of RNA PolII is so far the only other known substrate of CDKDs besides CDKA;1. Therefore, to assess the impact of CDKD;3 on transcription, we compare RNA-seq data obtained from rosette leaves of wild type, *cdkd;3-1* and *smg7-6* mutants as well as of *cdkd;3-1 smg7-6* double mutants. Interestingly, *cdkd;3-1* caused deregulation of only 114 genes compared to wild type with 105 genes upregulated and 9 genes downregulated (Figure 7a; Table S2). The vast majority of these genes (91/114) were also altered in *smg7-6* mutants, where in total 656 genes were deregulated. This is a striking result, as it indicates that CDKD3 targets almost exclusively genes also regulated by SMG7.

**Figure 7.**
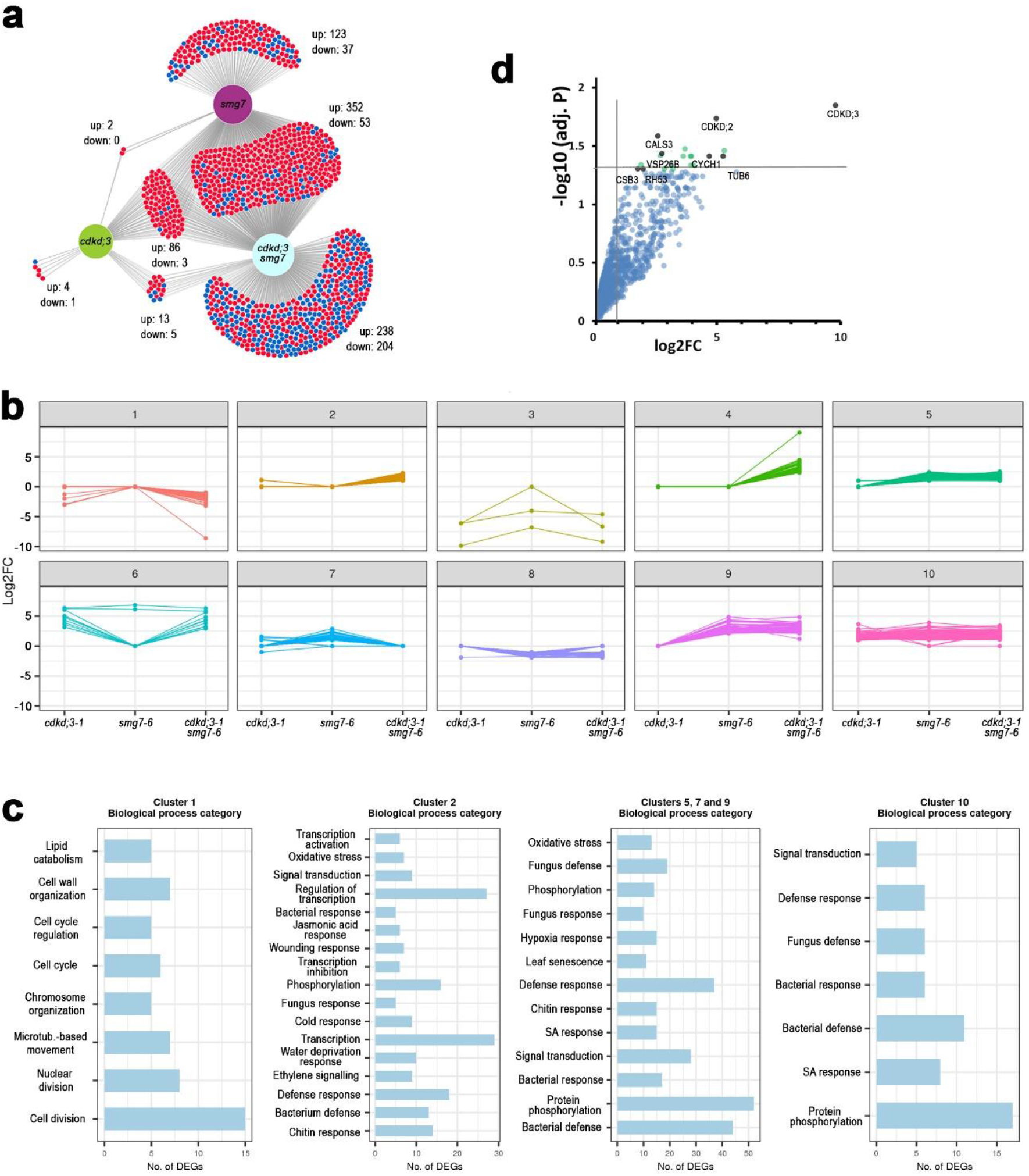
Transcriptomics and proteomics analysis of CDKD;3. (a) Venn diagram showing overlaps between differentially expressed genes relative to wild type in *cdkd;3-1, smg7-6* and *cdkd;3-1 smg7-6* mutants. Numbers of up- and down-regulated genes are indicated. (b) K-means clustering of DEGs. Clustering was performed on log2FC to build expression patterns in ten clusters for 1125 differentially expressed protein-coding genes. (c) Subsequent GO-term enrichment analysis of selected clusters. Genes in the clusters were analyzed with regard to their biological function by GO enrichment analysis (p-value < 0.05; Benjamini-Hochberg test). Most enriched GO categories are shown for representative categories of clusters. (d) Dot plot of proteins identified by LC-MS in pull-downs from wild type and CDKD;3:YFP expressing plants using GFP antibody. Proteins with the highest confidence of enrichment are indicated as black dots. Fold change (log2FC) and significance of the difference between wild type and CDKD;3:YFP samples are plotted.

To further analyze the expression datasets, we clustered genes based on their differential expression in different mutant combinations in ten clusters and performed gene ontology term (GO-term) enrichment analysis (Figure 7b,c). Genes upregulated in all three mutant combinations including *cdkd;3-1* (cluster 10) were almost exclusively enriched for GO-terms associated with pathogen defense response. Pathogen defense and oxidative stress were also enriched in clusters 5, 7 and 9, which include genes upregulated in *smg7-6* and *cdkd;3-1 smg7-6*, as well as in cluster 2 encompassing genes upregulated only in double mutants (Figure 7b,c). SMG7 controls pathogen response via suppressing plant intracellular immune receptors and its inactivation leads to a strong autoimmune response (Riehs-Kearnan *et al*., 2012, Gloggnitzer *et al*., 2014). This analysis indicates that even the relatively mild *smg7-6* allele with respect to NMD causes substantial upregulation of pathogen defense genes. CDKD;3 inactivation also leads to pathogen response, though much milder than in *smg7-6*. Moreover, *cdkd;3-1* further enhances pathogen and other stress responses when combined with *smg7-6* (cluster 2). The *cdkd;3-1 smg7-6* mutants also show a large portion of downregulated genes (Figure 7a). Their GO-term analysis revealed strong enrichment for genes involved in cell cycle and division (cluster 1, Figure 7b,c).

CDKD;3 might also phosphorylate additional substrates besides CDKA;1 and RNA PolII. To identify putative CDKD3;1 targets, we generated an Arabidopsis line expressing the CDKD;3::CDKD;3:YFP construct and performed immunoprecipitation coupled with mass spectrometry from floral bud protein extracts. The most significantly enriched proteins were CDKD;3, CDKD;2 and cyclin H, which reaffirmed the specificity of the immunoprecipitation protocol (Figure 7d). The other proteins with the highest confidence enrichment in the immunoprecipitated fraction were CALLOSE SYNTHASE 3 (CALS3; At5g13000), TUBULIN BETA-6 (TUBB6, At5g12250) and components of vesical trafficking systems COATOMER SUBUNIT BETA-3 (CSB3, At3g15980,) and VACUOLAR PROTEIN SORTING-ASSOCIATED PROTEIN 26B (VPS26B, At4g27690). Notably, the molecular functions of these proteins can be related to cytokinesis. While CDKD;3 resides most of the cell cycle in nuclei (Sofroni *et al*., 2020), it is released into the cytoplasm during the M-phase (Movie 7), and therefore, it has an opportunity to interact with these non-nuclear proteins.

## Discussion

Arabidopsis deficient in SMG7 exhibits an unusual meiotic phenotype that is characterized by a failure to complete meiosis and proceed to gametophytic development. Plants lacking SMG7 function arrest in meiotic anaphase II, whereas hypomorphic *smg7-6* mutants expressing SMG7 N-terminus form haploid nuclei that undergo multiple cycles of aberrant chromosome segregation without DNA replication (Riehs *et al*., 2008, Capitao *et al*., 2021). These additional meiotic cycles resemble meiosis II, where chromatid segregation also proceeds without DNA duplication. The key mechanism driving exit from the M-phase is CDK downregulation via anaphase promoting complex (APC/C) mediated degradation of M-type cyclins. This results in dephosphorylation of CDK substrates reverting molecular processes that led to the M-phase, and allowing for chromosome decondensation, spindle disassembly, formation of nuclei, and eventually cytokinesis (Potapova *et al*., 2006, Lopez-Aviles *et al*., 2009). Further decrease of CDK activity in G1 creates permissive conditions for licensing of origins of replication, which ensures the alternation of DNA replication and chromosome segregation. Therefore, the gradual downregulation of CDK activity provides order and directionality to molecular processes that occur during transition from M to G1. However, M-phase exit is not irreversible. Pharmacological manipulation of CDK activity during mitotic exit in human cells expressing non-degradable cyclin B could revert cells back to M-phase (Potapova *et al*., 2006). This principle is utilized in interkinesis between meiosis I and II when a partial inhibition of APC/C prevents full CDK inactivation and leads to reversion to M-phase prior to licensing of replication origins (Izawa *et al*., 2005, Cromer *et al*., 2012).

The Arabidopsis *smg7* phenotypes indicate insufficient downregulation of CDK activity at the end of meiosis. In support of this hypothesis, failure to fully activate APC/C and to downregulate CDK activity during meiotic exit in Drosophila and budding yeast have been reported to result in phenotypes similar to the ones described in *smg7* Arabidopsis (Page and Orr-Weaver, 1996, Chu *et al*., 2001, Wang *et al*., 2020). Thus, it is expected that mutations in genes that contribute to CDK activity would alleviate meiotic defects caused by SMG7 dysfunction. Indeed, we identified *cdkd;3-3* in the suppressor screen designed to uncover such mutations. The *cdkd;3-3* allele restores fertility of *smg7-6* plants by promoting orderly cytokinesis and tetrad formation, or, in PMCs that enter meiosis III, by delaying the onset of aberrant chromosome segregation after initiation of cytokinesis (Figure 2t,u). Because high CDK activity prevents transition to cytokinesis in plants (Sasabe *et al*., 2011), these results are consistent with the idea that CDKD3;1 is a CDK activator. CDKD;3 was reported to phosphorylate CDKA;1 and act as CDKA;1 activating kinase (Yamaguchi *et al*., 1998, Sofroni *et al*., 2020). However, we failed to mimic the *cdkd;3-3* effect on meiotic exit by the hypomorphic CDKA;1^VF^, which exhibits low CDK activity (Dissmeyer *et al*., 2009). This indicates that meiosis II is governed by another CDK, perhaps one of the CDKBs that were implicated to act together with B1 cyclins in microtubule organization during mitosis (Romeiro Motta *et al*., 2022).

The observation that CDKA;1^VF^ does not promote meiotic exit in *smg7-6* mutants opens up alternative scenarios, in which CDKD;3 affects meiotic exit through regulating other substrates than CDKs. Plant CDKDs are orthologous to human CDK7 kinases that are implicated in regulating transcription and RNA processing through phosphorylation of RNA Pol II (Joubes *et al*., 2000, Fisher, 2019). Whereas CDKD;3 has been reported to phosphorylate RNA Pol II *in vitro* (Yamaguchi *et al*., 1998, Shimotohno *et al*., 2003, Hajheidari *et al*., 2012), our transcriptomics analysis in *cdkd;3-1* mutant revealed only negligible impact on transcriptome likely due to functional redundancy between Arabidopsis CDKDs. Interestingly, the majority of upregulated transcripts were associated with pathogen response and almost perfectly overlapped with transcripts deregulated in *smg7-6* mutants. We observed that the combination of *cdkd;3-1* and *smg7-6* further enhanced upregulation of pathogen response and led to the downregulation of genes involved in cell cycle and division. Several pathogen signaling pathways were shown to inhibit the cell cycle (Qi and Zhang, 2019) indicating that upregulation of defense response could also contribute to the suppression of aberrant post-meiotic divisions in *smg7-6 cdkd;3* plants.

CDKD;3 may also phosphorylate additional proteins and affect the meiotic exit through direct regulation of proteins executing cell division. Remarkably, the top candidates we co-immunoprecipitated with CDKD;3 can be linked to cytokinesis. One such protein, TUB6 is part of the β-tubulin family that forms microtubules. Microtubules play a central role in mitosis and cytokinesis and phosphorylation of β-tubulin was proposed to regulate microtubule dynamics in human (Fourest-Lieuvin *et al*., 2006). Another candidate, CALS3, belongs to a family of callose synthases that mediate callose deposition in a wide range of cell types including meiocytes. Some members of the family have also been implicated in cytokines and microspore formation (Chen *et al*., 2009, De Storme and Geelen, 2013). We have also identified two proteins involved in endosomal trafficking important for the transport and turnover of plasma membrane proteins and cell wall biosynthetic enzymes. VPS26B is a subunit of the retromer complex that mediates protein recycling between Golgi apparatus, plasma membrane and vacuoles (Paez Valencia *et al*., 2016). CSB3 is a subunit of COPI coatomer complex responsible for Golgi vesicle trafficking, and its suppression results in aberrant cell plate formation during cytokinesis in tobacco cells (Ahn *et al*., 2015). To conclude, besides CDKA;1 and RNA Pol II, CDKD;3 may interact with and phosphorylate proteins involved in cytokinesis.

Our current work presents a more nuanced picture of the CDKD;3’s role in meiosis. In addition to dictating the simultaneous rather than sequential meiotic cytokinesis typical for dicots by inhibiting cell wall formation during interkinesis (Sofroni *et al*., 2020), CDKD;3 can also affect meiotic exit by influencing the rate of meiotic progression and onset of cytokinesis. These abilities are likely facilitated not only through regulation CDKA;1, but also additional protein substrates involved in cell division.

## Experimental procedures

### Plant material and growth conditions

*Arabidopsis thaliana* ecotype Columbia (Col-0) and mutant seeds were grown on soil in growth chambers at 21°C at 50–60% humidity with 16/8 h light/dark illumination. The following mutant lines were used in this study: *smg7-6* (Riehs-Kearnan *et al*., 2012), *tam-2* (Bulankova *et al*., 2010), *cdka;1-1* CDKA;1^VF^ (Dissmeyer *et al*., 2009), *cdkd;3-1, cdkd;2-1* and *cdkd;1-1* (Hajheidari *et al*., 2012). Mutations were assessed by PCR-genotyping using primers described in Table S3. Plants used for live-cell imaging were generated by crossing plants containing reporter constructs HTA10:RFP (Valuchova *et al*., 2020) and pRPS5A::TagRFP:TUB4 (Prusicki *et al*., 2019) with the parental *smg7-6* and the *EMS58-1* line.

### Generation of transgenic lines

For the complementation study, a PCR amplified CDKD;3 gene including 471 bp region upstream of the start codon and the 450bp downstream of stop codon was generated using the primers provided in Table S3. The amplicon was cloned into the vector pENTR™/ D-TOPO®Vector (Invitrogen), and transferred by Gateway LR reaction into the binary pGWB601 vector (Nakagawa *et al*., 2007). The construct was transformed by the floral dipping method into the *EMS58-1* line and T1 transformants were selected with Basta. The CDKD;3:YFP reporter line was generated the same Gateway cloning strategy by introducing the CDKD;3 gene fragment devoid of stop codon PCR amplified with primers indicated in Table S3 into the destination vector pGWB640 containing C terminus YFP tag. The resulting construct was transformed into wild-type Col-0 plants. Transformants were Basta selected and propagated up to the T3 generation.

### Assessment of plant fertility

Pollen viability was determined by Alexander staining (Alexander, 1969). The cell count plugin on FIJI (Schindelin *et al*., 2012) was used to count viable pollen. Silique length was measured by Images of main stems were captured using an Epson scanner and the siliques were measured using the FIJI Analyze/Measure function.

### Genetic mapping and complementation

*EMS58-1* seeds were obtained by the suppressor screening in *smg7-6* background described in (Capitao *et al*., 2021). Mutations associated with improved fertility were identified by whole genome sequencing of pooled fertile plants using ArtMAP (Javorka *et al*., 2019). The identified *cdkd;3-3* mutation was genotyped by the High-Resolution Melting (HRM) qPCR-based method using primers described in Table S3 using Lightcycler96 thermocycler (Roche) using the inbuilt HRM profile.

### Cytology

Staining of PMCs in whole anthers was performed as previously described (Capitao *et al*., 2021) using 2ug/ml DAPI stock solution for DNA staining or 0.1% (v/v) solution of SR2200 (Renchem) for callose staining. Images were acquired by LSM700 confocal microscope (1024p resolution, Emission/Absorption wavelengths of 405/435, 1AU, 35% laser power).

### Live cell imaging

Live imaging was performed as previously described (Valuchova *et al*., 2020). Floral buds 0.3 to 0.7 mm long were selected from the main inflorescence. The reproductive organs were exposed by removing the sepals and placed into glass capillaries (size 4, Zeiss) containing ½ MS medium (5% sucrose, pH 5.8) with 1% low melting point agarose (Sigma Aldrich). The inflorescence embedded in solidified MS was pushed out of the capillary and then placed into the capillary holder for the Z.1 ZEISS light-sheet microscope. After the holder is inserted in the microscope chamber, the remaining space is filled with liquid ½ MS medium (5% sucrose, pH 5.8). Images were taken every 5 minutes with a 10x objective (Detection optics 10x/0.5), single illumination (Illumination Optics 10x/0.2), 561 nm laser (15% intensity). The large raw data files were processed by ZEN Blue software (Zeiss).

### Transcriptome analysis

Leaf tissue was collected from three weeks-old plants and total RNA was extracted using RNA Blue (Top-Bio), following the manufacturer’s protocol. For each sample, 10 μg of total RNA was treated with DNase I (Roche). Strand-specific libraries were prepared from ribosomal RNA-depleted total RNA using ScriptSeq complete kit for plant leaf (Epicentre; BPL1224) following manufacturer’s protocol. After assessing the quality and quantity of RNASeq libraries by fragment analyzer, libraries were 125/150 bp paired-end sequenced on Illumina HiSeqv4/NextSeq. Low-quality reads and adapters were trimmed from the total read pairs using cutadapt v2.5. Then, the RNA-Seq data was aligned on reference AtRTD2 transcript dataset (Zhang *et al*., 2017) and simultaneously quantified with quasi-mapping-based algorithm using Salmon v1.5 (Patro *et al*., 2017). The downstream data normalization and differential expression analysis was done using 3D RNA-Seq (Calixto *et al*., 2018). A gene was significantly differentially expressed (DE) in a contrast group if it had adjusted p-value < 0.05 (Benjamini-Hochberg (BH)-corrected (Benjamini and Yekutieli, 2001), and absolute log2FC ≥ 1. The transcript overlaps among datasets were visualized by a custom Venn diagram tool available at https://divenn.tch.harvard.edu/. Further, functional annotation and gene ontology (GO) enrichment was done online via DAVID bioinformatic database (https://david.ncifcrf.gov/summary.jsp;(Sherman *et al*., 2022). Next, to categorize the DE genes with similar expression patterns, k-means clustering was performed and plots were generated using R packages.

### Immunoprecipitation and mass spectrometry

Meiotic buds from ∼150 inflorescences from CDKD;3:YFP and wild type control were grinded in liquid nitrogen and processed using the ChromoTek GFP-Trap® Magnetic Agarose kit (Chromotek). Briefly, the sample was, mixed with 200 µl of RIPA buffer supplemented with 2.5 μl of DNAse I (NEB), 10 μl of 50 mM MgCl_2_ and 3.5 μl of protease and phosphatase inhibitor cocktail (Halt), and incubated on ice for 45 min. Lysates were sedimented by centrifugation and the supernatant was transferred to a new 1.5 ml tube containing 450 μl ice-cold dilution buffer included in the kit and 6.5 μl of protease and phosphatase inhibitor cocktail. 200 μg of the extracted proteins were added to 25 μl of equilibrated magnetic agarose beads (Chromotek) and mixed in a rotating shaker for 45 min at 4 degrees. The supernatant was mixed with equilibrated ChromoTek GFP-Trap® Magnetic Agarose beads GFP-Trap MA beads and incubated for 90 min at 4°C. After washing with the dilution buffer, the bead bound protein complexes were digested directly on beads by addition of 0.5 μg (1μg/μl) of trypsin (sequencing grade, Promega) in 50mM NaHCO3 buffer and incubated at 37°C for 18 h. Resulting peptides were extracted into LC-MS vials by 2.5% formic acid and acetonitrile with addition of polyethylene glycol (20,000; final concentration 0.001%) and concentrated in a SpeedVac concentrator (Thermo Fisher Scientific).

LC-MS/MS analyses of all peptide mixtures were done using Ultimate 3000 RSLCnano system connected to Orbitrap Fusion Lumos Tribrid mass spectrometer (Thermo Fisher Scientific). Prior to LC separation, tryptic digests were online concentrated and desalted using trapping column (100 μm × 30 mm, 3.5μm particles, X-Bridge BEH 130 C18 sorbent (Waters, Milford, MA, USA; temperature of 40 °C) After washing of trapping column with 0.1% FA, the peptides were eluted (flow rate - 500 nl/min) from the trapping column onto an analytical column (Acclaim Pepmap100 C18, 3 µm particles, 75 μm × 500 mm; column compartment temperature of 40 °C, Thermo Fisher Scientific) by 135 min nonlinear gradient program (1-56% of mobile phase B; mobile phase A: 0.1% FA in water; mobile phase B: 0.1% FA in 80% ACN). The analytical column outlet was directly connected to the Digital PicoView 550 (New Objective) ion source with sheath gas option and SilicaTip emitter (New Objective; FS360-20-15-N-20-C12) utilization. ABIRD (Active Background Ion Reduction Device, ESI Source Solutions) was installed.

MS data were acquired in a data-dependent strategy with defined number of scans based on precursor abundance with survey scan (*m/z* 350-2000). The resolution of the survey scan was 120000 (at *m/z* 200) with a target value of 4×10^5^ ions and maximum injection time of 100 ms. HCD MS/MS (30% relative fragmentation energy, normal mass range) spectra were acquired with a target value of 5.0×10^4^ and resolution of 15 000 (at *m/z* 200). The maximum injection time for MS/MS was 22 ms. Dynamic exclusion was enabled for 30 s after one MS/MS spectra acquisition. The isolation window for MS/MS fragmentation was set to 1.2 *m/z*.

The analysis of the mass spectrometric RAW data files was carried out using the MaxQuant software (version 1.6.2.0) using default settings unless otherwise noted. MS/MS ion searches were done against modified cRAP database (based on http://www.thegpm.org/crap) containing protein contaminants like keratin, trypsin etc., and UniProtKB protein database for *Arabidopsis thaliana* (ftp://ftp.uniprot.org/pub/databases/uniprot/current_release/knowledgebase/reference_proteomes/Eukaryota/UP000006548_3702.fasta.gz; downloaded 05.2018, version 2018/05, number of protein sequences: 27,582). Oxidation of methionine and proline, deamidation (N, Q) and acetylation (protein N-terminus) as optional modification, and trypsin/P enzyme with 2 allowed missed cleavages were set. Peptides and proteins with FDR threshold <0.01 and proteins having at least one unique or razor peptide were considered only. Match between runs was set among all analysed samples. Protein abundance was assessed using protein intensities calculated by MaxQuant. Protein intensities reported in proteinGroups.txt file (output of MaxQuant) were further processed using the software container environment (https://github.com/OmicsWorkflows), version 3.7.2a.

## Acknowledgement

We acknowledge the support from CEITEC MU Core facilities Plant Sciences, Proteomics, and CELLIM, supported by the Czech-Bioimaging (No. LM2018129) and CIISB (No. LM2018127) infrastructure project projects funded by MEYES CZ. The genome sequencing was performed by the Next Generation Sequencing Facility at Vienna BioCenter Core Facilities (VBCF), member of the Vienna BioCenter (VBC), Austria. This work was supported by Ministry of Education, Youth, and Sports of the Czech Republic, the European Regional Development Fund-Project ‘REMAP’ (No. CZ.02.1.01/0.0/0.0/15_003/0000479 to K.R.), by the Czech Science Foundation (grant 21-25163J to K.R.) and German Research Foundation (grant 452003411 to A.S.)

---

**Movie 1**. Time-lapse microscopy of meiotic divisions in wild type PMCs from diakinesis till telophase II in 5 min increments. Chromatin was labeled with HTA10:TagRFP reporter. Scale bar = 20 µm.

**Movie 2**. Time-lapse microscopy of meiotic divisions in *smg7-6* PMCs from diakinesis till telophase III in 5 min increments. Chromatin was labeled with HTA10:TagRFP reporter. Scale bar = 20 µm.

**Movie 3**. Time-lapse microscopy of meiotic divisions in *EMS58-1* PMCs from diakinesis till telophase III in 5 min increments. Chromatin was labeled with HTA10:TagRFP reporter. Scale bar = 20 µm.

**Movie 4**. Time-lapse microscopy of meiotic divisions in wild type PMCs from interkinesis till the tetrad stage in 5 min increments. Microtubules were labeled with GFP:TUA5 reporter. Scale bar = 20 µm.

**Movie 5**. Time-lapse microscopy of meiotic divisions in *smg7-6* PMCs from interkinesis, through meiosis II and meiosis III till the polyad stage in 5 min increments. Microtubules were labeled with GFP:TUA5 reporter. Scale bar = 20 µm.

**Movie 6**. Time-lapse microscopy of meiotic divisions in *EMS58-1* PMCs from interkinesis, through meiosis II and meiosis III till the polyad stage in 5 min increments. Note that some PMCs do not proceed in meiosis III and form regular tetrads. Microtubules were labeled with GFP:TUA5 reporter. Scale bar = 20 µm.

**Movie 7**. Time-lapse microscopy of CDKD;3 localization in PMCs from late prophase I till telophase II s in 5 min increments. CDKD;3 and chromatin were visualized using CDKD;3:YFP (green) and HTA10:TagRFP (magenta) reporters, respectively.

## Supplementary figures

**Figure S1.**
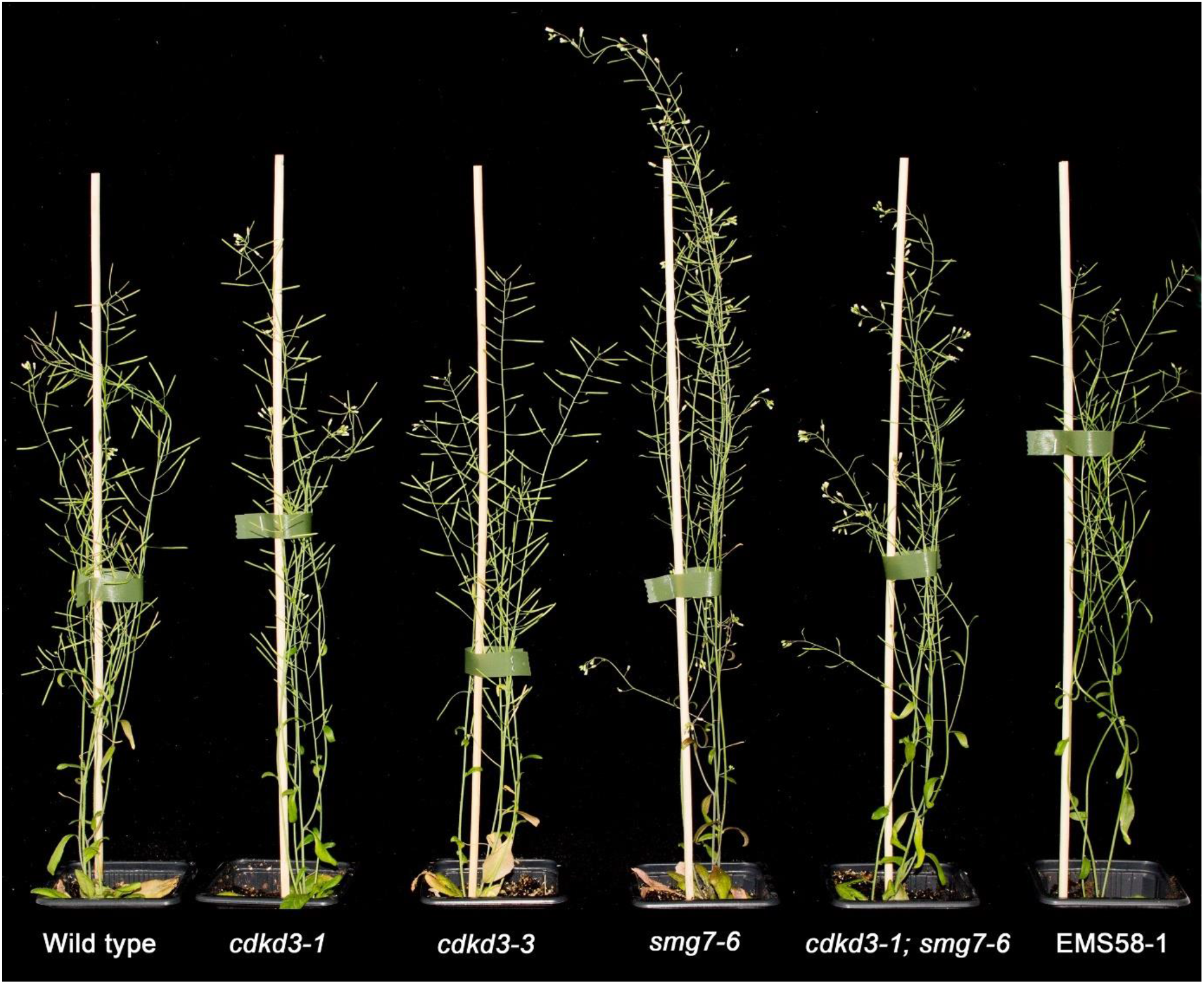
Arabidopsis *cdkd* and *smg7-6* mutants. Approximately 6-weks old plants are shown. Longer inflorescence bolt in the *smg7-6* mutant is caused by reduced fertility.

**Figure S2.**
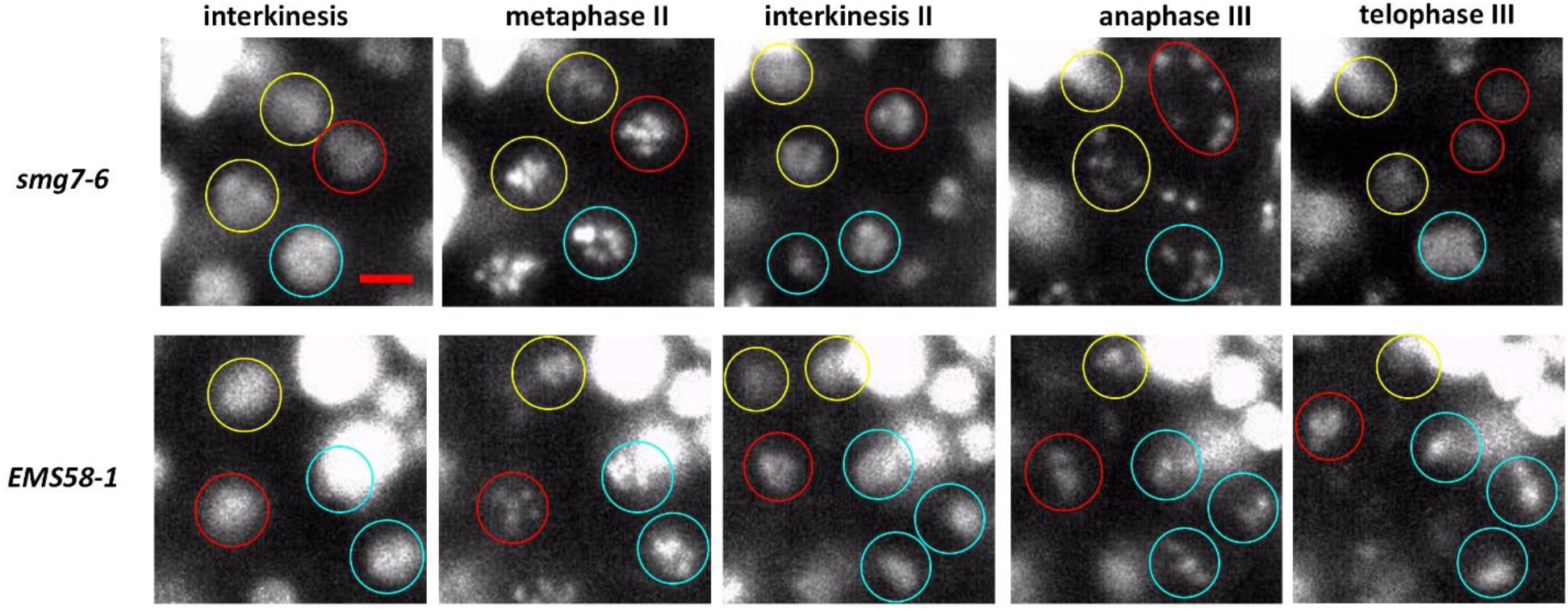
Time-lapse series of *smg7-6* and EMS58-1 meiotic nuclei undergoing meiosis II and meiosis III. Nuclei derived from the same PMC are encircled with identical color. Note that chromatid tends to spread over larger area during anaphase III in *smg7-6* mutants. Chromatin is visualized with HTA10:TagRFP marker. Scale bar = 5 µm.

**Figure S3.**
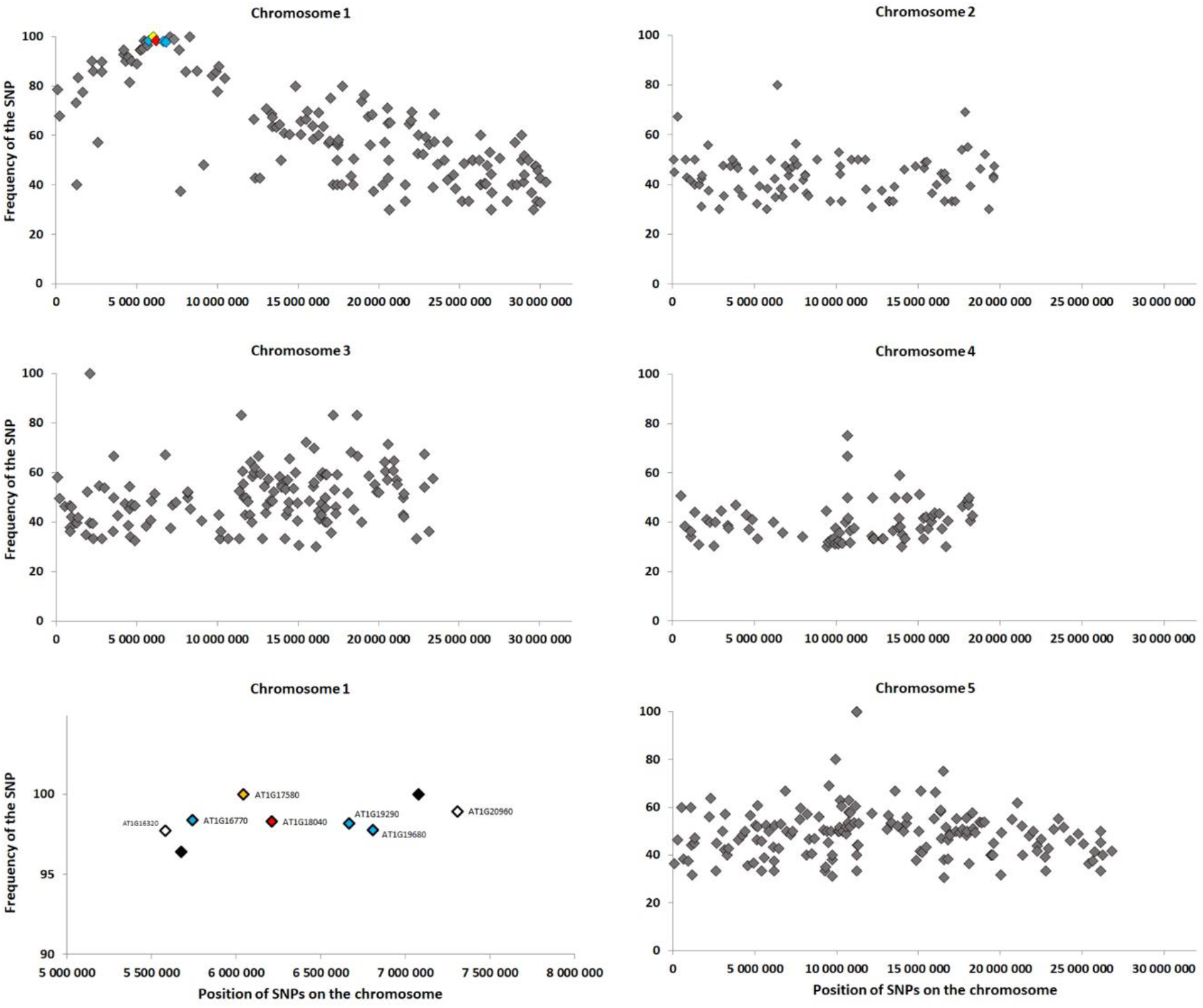
Mapping mutations associated with the rescued fertility in *EMS58-1* by whole genome sequencing. The causative mutation is recessive and is, therefore, expected to fully associate with the rescued phenotype. The charts indicate positions of *de novo* mutations on individual Arabidopsis chromosomes (x axis) and their frequency in population of B2 plants exhibiting increased fertility. Colored diamonds on chromosome 1 mark *de novo* mutations with the highest association. This region is zoomed in the lower left graph with indicated AGI codes for the mutated genes.

**Figure S4.**
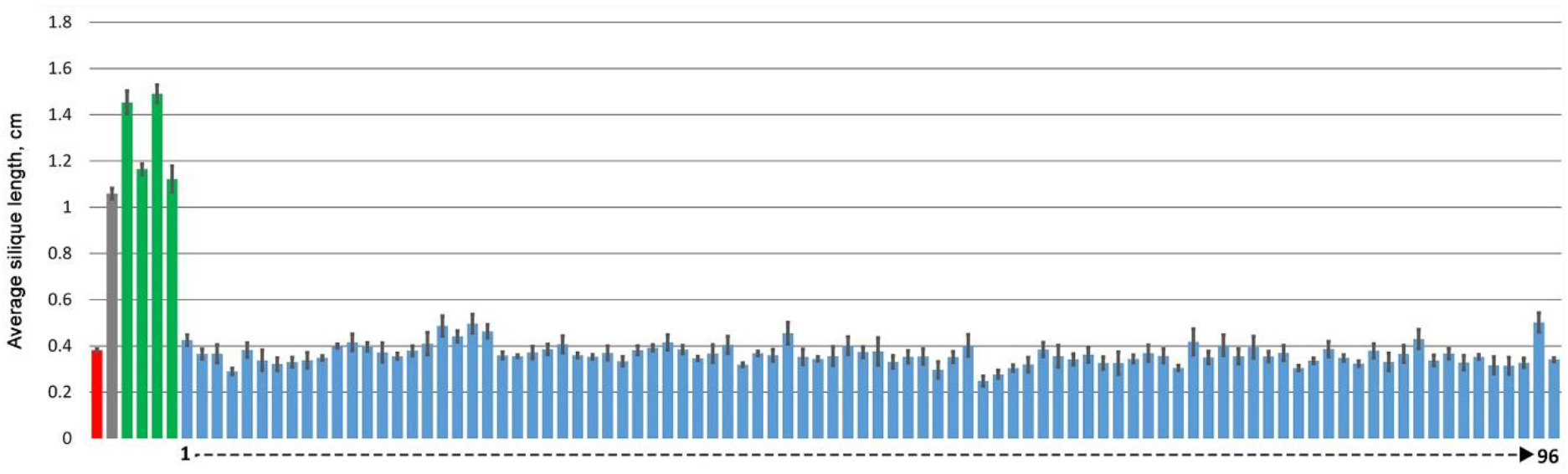
Genetic complementation of *EMS58-1* plants with the *CDKD;3* genic construct causes infertility in transformed T1 plants. The chart shows average silique length in individual T1 transformants (error bars indicate SD; n= 25). Data for *smg7-6* are shown in red, for EMS58-1 in gray, for complemented T1 transformants in blue and for non-complemented transformants in green.

**Figure S5.**
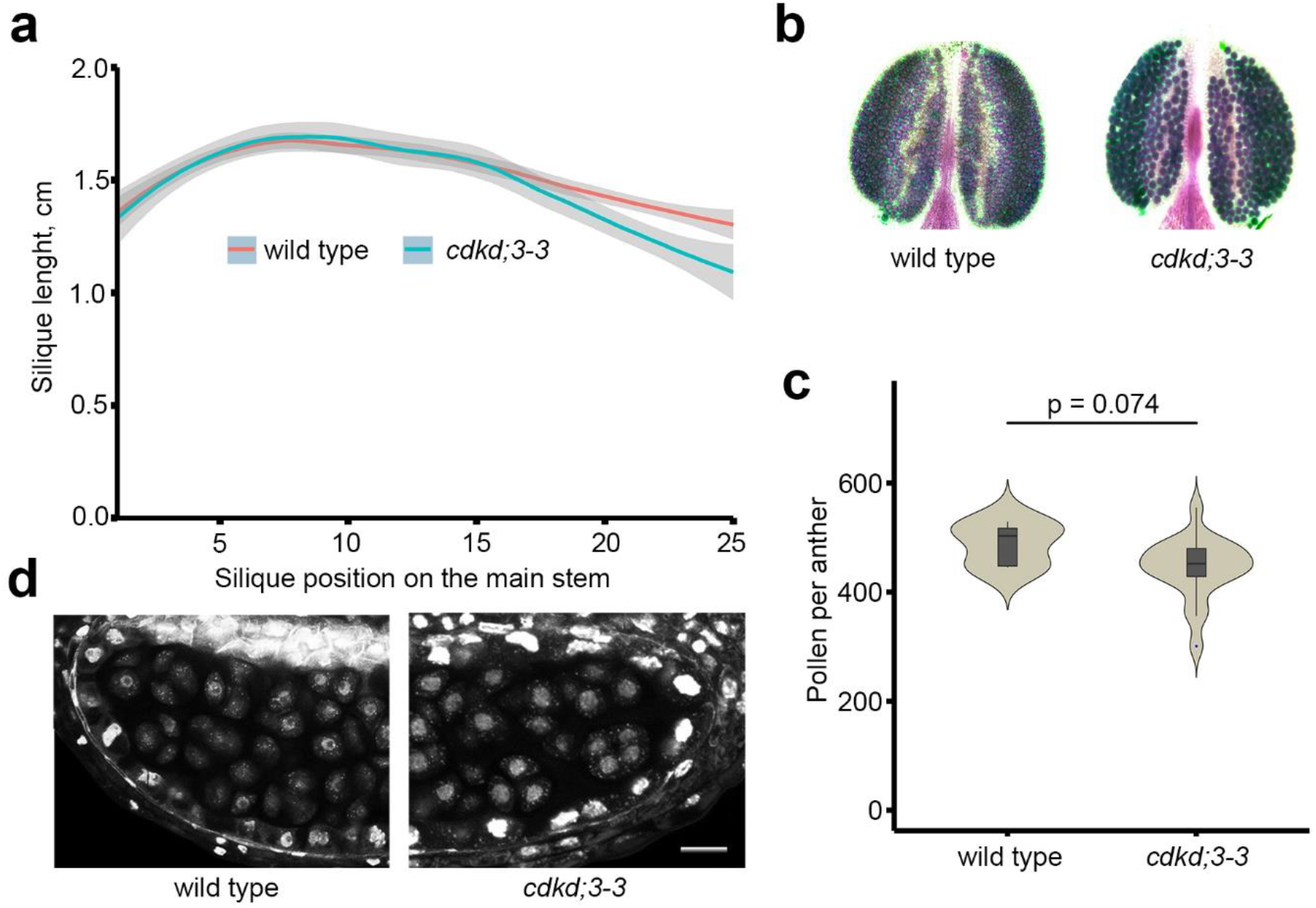
Fertility of *cdkd;3-3* mutants. (a) Quantification of silique length along the main inflorescence bolt in wild type (n = 10) and cdkd;3-3 (n = 5). The trend lines of the data were plotted by LOESS smooth function (colored line; shaded area represents 95% confidence intervals). (b) Anthers from wild type and *cdkd;3-3* plants after Alexander staining. (c) Violin plots showing viable pollen per anther. Significance of the difference is indicated (two tailed t-test; wild type n = 20; *cdkd;3-3* = 5). (d) Tetrads in anther lobes stained with DAPI.

**Figure S6.**
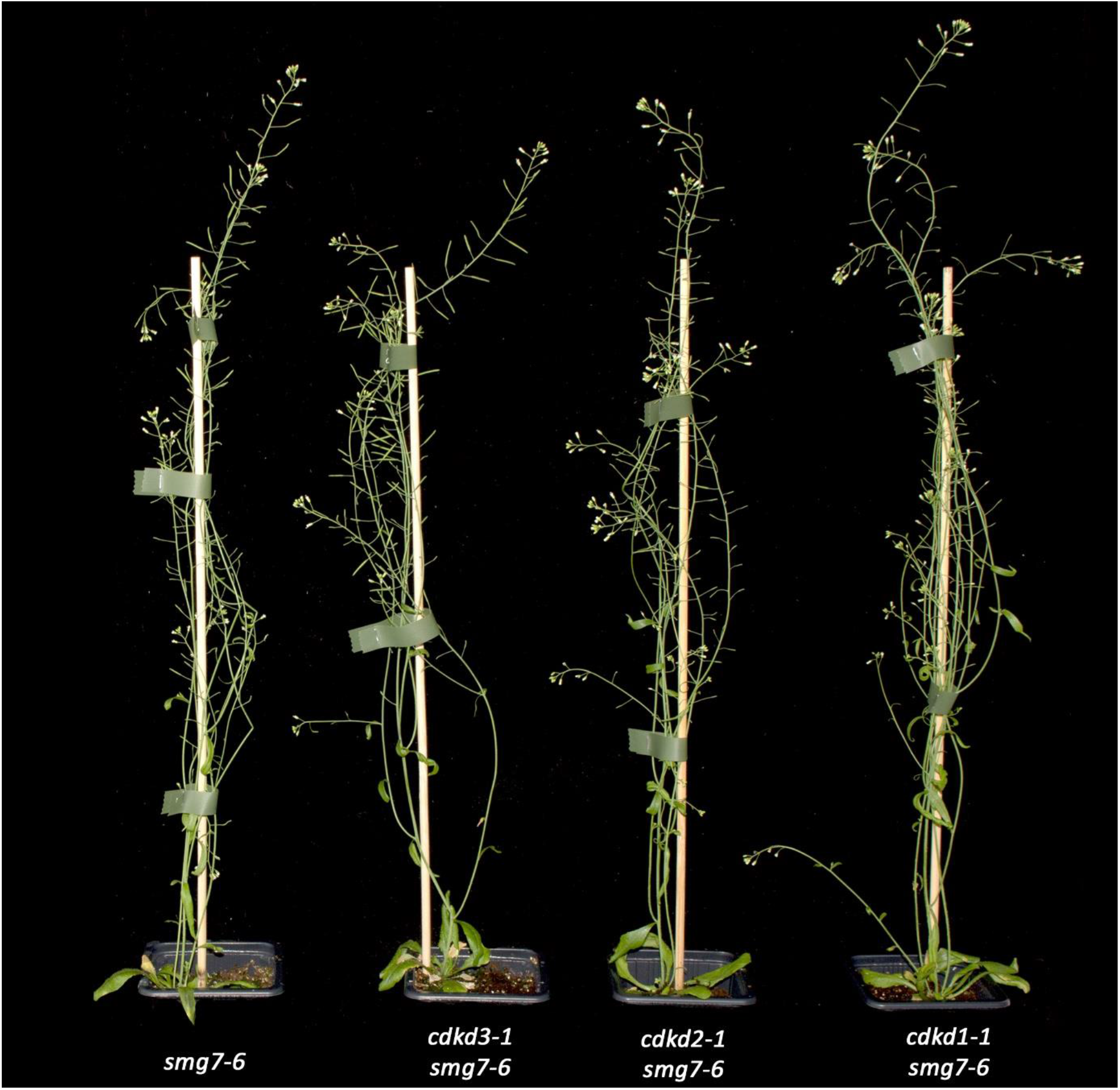
Approximately six weeks old mutant plants.

## Notes

### Competing Interest Statement

The authors have declared no competing interest.

## Literature

Ahn, H.K., Kang, Y.W., Lim, H.M., Hwang, I. and Pai, H.S. (2015) Physiological Functions of the COPI Complex in Higher Plants. Mol Cells, 38, 866–875.

Alexander, M.P. (1969) Differential staining of aborted and nonaborted pollen. Stain Technology, 44, 117–122.

Azumi, Y., Liu, D., Zhao, D., Li, W., Wang, G., Hu, Y. and Ma, H. (2002) Homolog interaction during meiotic prophase I in Arabidopsis requires the SOLO DANCERS gene encoding a novel cyclin-like protein. Embo J, 21, 3081–3095.

Benjamini, Y. and Yekutieli, D. (2001) The control of the false discovery rate in multiple testing under dependency. Ann Stat, 29, 1165–1188.

Bulankova, P., Akimcheva, S., Fellner, N. and Riha, K. (2013) Identification of Arabidopsis meiotic cyclins reveals functional diversification among plant cyclin genes. PLoS Genet, 9, e1003508.

Bulankova, P., Riehs-Kearnan, N., Nowack, M.K., Schnittger, A. and Riha, K. (2010) Meiotic progression in Arabidopsis is governed by complex regulatory interactions between SMG7, TDM1, and the meiosis I-specific cyclin TAM. Plant Cell, 22, 3791–3803.

Cairo, A., Vargova, A., Shukla, N., Capitao, C., Mikulkova, P., Valuchova, S., Pecinkova, J., Bulankova, P. and Riha, K. (2022) Meiotic exit in Arabidopsis is driven by P-body-mediated inhibition of translation. Science, 377, 629–634.

Calixto, C.P.G., Guo, W., James, A.B., Tzioutziou, N.A., Entizne, J.C., Panter, P.E., Knight, H., Nimmo, H.G., Zhang, R. and Brown, J.W.S. (2018) Rapid and Dynamic Alternative Splicing Impacts the Arabidopsis Cold Response Transcriptome. Plant Cell, 30, 1424–1444.

Capitao, C., Tanasa, S., Fulnecek, J., Raxwal, V.K., Akimcheva, S., Bulankova, P., Mikulkova, P., Crhak Khaitova, L., Kalidass, M., Lermontova, I., Mittelsten Scheid, O. and Riha, K. (2021) A CENH3 mutation promotes meiotic exit and restores fertility in SMG7-deficient Arabidopsis. PLoS Genet, 17, e1009779.

Cromer, L., Heyman, J., Touati, S., Harashima, H., Araou, E., Girard, C., Horlow, C., Wassmann, K., Schnittger, A., De Veylder, L. and Mercier, R. (2012) OSD1 promotes meiotic progression via APC/C inhibition and forms a regulatory network with TDM and CYCA1;2/TAM. PLoS Genet, 8, e1002865.

De Storme, N. and Geelen, D. (2013) Cytokinesis in plant male meiosis. Plant Signal Behav, 8, e23394.

Dissmeyer, N., Nowack, M.K., Pusch, S., Stals, H., Inze, D., Grini, P.E. and Schnittger, A. (2007) T-loop phosphorylation of Arabidopsis CDKA;1 is required for its function and can be partially substituted by an aspartate residue. Plant Cell, 19, 972–985.

Dissmeyer, N., Weimer, A.K., Pusch, S., De Schutter, K., Kamei, C.L., Nowack, M.K., Novak, B., Duan, G.L., Zhu, Y.G., De Veylder, L. and Schnittger, A. (2009) Control of cell proliferation, organ growth, and DNA damage response operate independently of dephosphorylation of the Arabidopsis Cdk1 homolog CDKA;1. Plant Cell, 21, 3641–3654.

Fisher, R.P. (2019) Cdk7: a kinase at the core of transcription and in the crosshairs of cancer drug discovery. Transcription, 10, 47–56.

Fourest-Lieuvin, A., Peris, L., Gache, V., Garcia-Saez, I., Juillan-Binard, C., Lantez, V. and Job, D. (2006) Microtubule regulation in mitosis: tubulin phosphorylation by the cyclin-dependent kinase Cdk1. Mol Biol Cell, 17, 1041–1050.

Francis, K.E., Lam, S.Y. and Copenhaver, G.P. (2006) Separation of Arabidopsis pollen tetrads is regulated by QUARTET1, a pectin methylesterase gene. Plant Physiol, 142, 1004–1013.

Gloggnitzer, J., Akimcheva, S., Srinivasan, A., Kusenda, B., Riehs, N., Stampfl, H., Bautor, J., Dekrout, B., Jonak, C., Jimenez-Gomez, J.M., Parker, J.E. and Riha, K. (2014) Nonsense-mediated mRNA decay modulates immune receptor levels to regulate plant antibacterial defense. Cell Host Microbe, 16, 376–390.

Hajheidari, M., Farrona, S., Huettel, B., Koncz, Z. and Koncz, C. (2012) CDKF;1 and CDKD protein kinases regulate phosphorylation of serine residues in the C-terminal domain of Arabidopsis RNA polymerase II. Plant Cell, 24, 1626–1642.

Harashima, H., Shinmyo, A. and Sekine, M. (2007) Phosphorylation of threonine 161 in plant cyclin-dependent kinase A is required for cell division by activation of its associated kinase. Plant J, 52, 435–448.

Chen, X.Y., Liu, L., Lee, E., Han, X., Rim, Y., Chu, H., Kim, S.W., Sack, F. and Kim, J.Y. (2009) The Arabidopsis callose synthase gene GSL8 is required for cytokinesis and cell patterning. Plant Physiol, 150, 105–113.

Chu, T., Henrion, G., Haegeli, V. and Strickland, S. (2001) Cortex, a Drosophila gene required to complete oocyte meiosis, is a member of the Cdc20/fizzy protein family. Genesis, 29, 141–152.

Izawa, D., Goto, M., Yamashita, A., Yamano, H. and Yamamoto, M. (2005) Fission yeast Mes1p ensures the onset of meiosis II by blocking degradation of cyclin Cdc13p. Nature, 434, 529–533.

Javorka, P., Raxwal, V.K., Najvarek, J. and Riha, K. (2019) artMAP: A user-friendly tool for mapping ethyl methanesulfonate-induced mutations in Arabidopsis. Plant Direct, 3, e00146.

Joubes, J., Chevalier, C., Dudits, D., Heberle-Bors, E., Inze, D., Umeda, M. and Renaudin, J.P. (2000) CDK-related protein kinases in plants. Plant Mol Biol, 43, 607–620.

Lopez-Aviles, S., Kapuy, O., Novak, B. and Uhlmann, F. (2009) Irreversibility of mitotic exit is the consequence of systems-level feedback. Nature, 459, 592–595.

Nakagawa, T., Suzuki, T., Murata, S., Nakamura, S., Hino, T., Maeo, K., Tabata, R., Kawai, T., Tanaka, K., Niwa, Y., Watanabe, Y., Nakamura, K., Kimura, T. and Ishiguro, S. (2007) Improved Gateway binary vectors: high-performance vectors for creation of fusion constructs in transgenic analysis of plants. Biosci Biotechnol Biochem, 71, 2095–2100.

Paez Valencia, J., Goodman, K. and Otegui, M.S. (2016) Endocytosis and Endosomal Trafficking in Plants. Annu Rev Plant Biol, 67, 309–335.

Page, A.W. and Orr-Weaver, T.L. (1996) The Drosophila genes grauzone and cortex are necessary for proper female meiosis. J Cell Sci, 109 (Pt 7), 1707–1715.

Parry, D.H. and O’Farrell, P.H. (2001) The schedule of destruction of three mitotic cyclins can dictate the timing of events during exit from mitosis. Curr Biol, 11, 671–683.

Patro, R., Duggal, G., Love, M.I., Irizarry, R.A. and Kingsford, C. (2017) Salmon provides fast and bias-aware quantification of transcript expression. Nat Methods, 14, 417–419.

Potapova, T.A., Daum, J.R., Pittman, B.D., Hudson, J.R., Jones, T.N., Satinover, D.L., Stukenberg, P.T. and Gorbsky, G.J. (2006) The reversibility of mitotic exit in vertebrate cells. Nature, 440, 954–958.

Prusicki, M.A., Keizer, E.M., van Rosmalen, R.P., Komaki, S., Seifert, F., Muller, K., Wijnker, E., Fleck, C. and Schnittger, A. (2019) Live cell imaging of meiosis in Arabidopsis thaliana. Elife, 8.

Qi, F. and Zhang, F. (2019) Cell Cycle Regulation in the Plant Response to Stress. Front Plant Sci, 10, 1765.

Riehs-Kearnan, N., Gloggnitzer, J., Dekrout, B., Jonak, C. and Riha, K. (2012) Aberrant growth and lethality of Arabidopsis deficient in nonsense-mediated RNA decay factors is caused by autoimmune-like response. Nucleic Acids Res, 40, 5615–5624.

Riehs, N., Akimcheva, S., Puizina, J., Bulankova, P., Idol, R.A., Siroky, J., Schleiffer, A., Schweizer, D., Shippen, D.E. and Riha, K. (2008) Arabidopsis SMG7 protein is required for exit from meiosis. J Cell Sci, 121, 2208–2216.

Romeiro Motta, M., Zhao, X., Pastuglia, M., Belcram, K., Roodbarkelari, F., Komaki, M., Harashima, H., Komaki, S., Kumar, M., Bulankova, P., Heese, M., Riha, K., Bouchez, D. and Schnittger, A. (2022) B1-type cyclins control microtubule organization during cell division in Arabidopsis. EMBO Rep, 23, e53995.

Sasabe, M., Boudolf, V., De Veylder, L., Inze, D., Genschik, P. and Machida, Y. (2011) Phosphorylation of a mitotic kinesin-like protein and a MAPKKK by cyclin-dependent kinases (CDKs) is involved in the transition to cytokinesis in plants. Proc Natl Acad Sci U S A, 108, 17844–17849.

Sherman, B.T., Hao, M., Qiu, J., Jiao, X., Baseler, M.W., Lane, H.C., Imamichi, T. and Chang, W. (2022) DAVID: a web server for functional enrichment analysis and functional annotation of gene lists (2021 update). Nucleic Acids Res.

Shimotohno, A., Aki, S.S., Takahashi, N. and Umeda, M. (2021) Regulation of the Plant Cell Cycle in Response to Hormones and the Environment. Annu Rev Plant Biol, 72, 273–296.

Shimotohno, A., Matsubayashi, S., Yamaguchi, M., Uchimiya, H. and Umeda, M. (2003) Differential phosphorylation activities of CDK-activating kinases in Arabidopsis thaliana. FEBS Lett, 534, 69–74.

Schindelin, J., Arganda-Carreras, I., Frise, E., Kaynig, V., Longair, M., Pietzsch, T., Preibisch, S., Rueden, C., Saalfeld, S., Schmid, B., Tinevez, J.Y., White, D.J., Hartenstein, V., Eliceiri, K., Tomancak, P. and Cardona, A. (2012) Fiji: an open-source platform for biological-image analysis. Nat Methods, 9, 676–682.

Sofroni, K., Takatsuka, H., Yang, C., Dissmeyer, N., Komaki, S., Hamamura, Y., Bottger, L., Umeda, M. and Schnittger, A. (2020) CDKD-dependent activation of CDKA;1 controls microtubule dynamics and cytokinesis during meiosis. J Cell Biol, 219.

Takatsuka, H., Umeda-Hara, C. and Umeda, M. (2015) Cyclin-dependent kinase-activating kinases CDKD;1 and CDKD;3 are essential for preserving mitotic activity in Arabidopsis thaliana. Plant J, 82, 1004–1017.

Umeda, M., Shimotohno, A. and Yamaguchi, M. (2005) Control of cell division and transcription by cyclin-dependent kinase-activating kinases in plants. Plant Cell Physiol, 46, 1437–1442.

Valuchova, S., Mikulkova, P., Pecinkova, J., Klimova, J., Krumnikl, M., Bainar, P., Heckmann, S., Tomancak, P. and Riha, K. (2020) Imaging plant germline differentiation within Arabidopsis flowers by light sheet microscopy. Elife, 9.

Vandepoele, K., Raes, J., De Veylder, L., Rouze, P., Rombauts, S. and Inze, D. (2002) Genome-wide analysis of core cell cycle genes in Arabidopsis. Plant Cell, 14, 903–916.

Wang, F., Zhang, R., Feng, W., Tsuchiya, D., Ballew, O., Li, J., Denic, V. and Lacefield, S. (2020) Autophagy of an Amyloid-like Translational Repressor Regulates Meiotic Exit. Dev Cell, 52, 141–151 e145.

Wang, G., Kong, H., Sun, Y., Zhang, X., Zhang, W., Altman, N., DePamphilis, C.W. and Ma, H. (2004a) Genome-wide analysis of the cyclin family in Arabidopsis and comparative phylogenetic analysis of plant cyclin-like proteins. Plant Physiol, 135, 1084–1099.

Wang, Y., Magnard, J.L., McCormick, S. and Yang, M. (2004b) Progression through meiosis I and meiosis II in Arabidopsis anthers is regulated by an A-type cyclin predominately expressed in prophase I. Plant Physiol, 136, 4127–4135.

Wijnker, E., Harashima, H., Muller, K., Parra-Nunez, P., de Snoo, C.B., van de Belt, J., Dissmeyer, N., Bayer, M., Pradillo, M. and Schnittger, A. (2019) The Cdk1/Cdk2 homolog CDKA;1 controls the recombination landscape in Arabidopsis. Proc Natl Acad Sci U S A, 116, 12534–12539.

Yamaguchi, M., Umeda, M. and Uchimiya, H. (1998) A rice homolog of Cdk7/MO15 phosphorylates both cyclin-dependent protein kinases and the carboxy-terminal domain of RNA polymerase II. Plant J, 16, 613–619.

Yang, C., Sofroni, K., Wijnker, E., Hamamura, Y., Carstens, L., Harashima, H., Stolze, S.C., Vezon, D., Chelysheva, L., Orban-Nemeth, Z., Pochon, G., Nakagami, H., Schlogelhofer, P., Grelon, M. and Schnittger, A. (2020) The Arabidopsis Cdk1/Cdk2 homolog CDKA;1 controls chromosome axis assembly during plant meiosis. EMBO J, 39, e101625.

Zhang, R., Calixto, C.P.G., Marquez, Y., Venhuizen, P., Tzioutziou, N.A., Guo, W., Spensley, M., Entizne, J.C., Lewandowska, D., Have, S.T., Frey, N.F., Hirt, H., James, A.B., Nimmo, H.G., Barta, A., Kalyna, M. and Brown, J.W.S. (2017) A high quality Arabidopsis transcriptome for accurate transcript-level analysis of alternative splicing. Nucleic Acids Research.

